# Physiological Arousal as a Predominant Source of Individual Differences in Functional Brain Networks

**DOI:** 10.64898/2025.12.18.694946

**Authors:** Justine A. Hill, Tianye Zhai, Cole Korponay, Julia C. Welsh, Betty Jo Salmeron, Thomas J. Ross, Yihong Yang, Blaise deB. Frederick, Amy C. Janes

**Affiliations:** National Institute on Drug Abuse Intramural Research Program, Biomedical Research Center 251 Bayview Blvd. 7A Baltimore, MD 21224; McLean Hospital, Belmont, MA; Harvard Medical School, Boston, MA

## Abstract

Individual differences in brain network function and organization are promising targets for fMRI-based biomarkers in precision psychiatry, yet the sources of individual variability remain poorly understood. We show that arousal, assessed by systemic low-frequency oscillation (sLFO) amplitude in the fMRI signal, accounts for a substantial portion of variance in brain network properties (e.g.: R= 0.70 for default mode network (DMN)–dorsal attention connectivity; R= –0.63 for DMN dynamics). These relationships replicated across sessions, across independent samples, and when assessed with traditional arousal indices. Critically, associations persisted after sLFO denoising, indicating a genuine brain-physiology relationship rather than hemodynamic artifact. Pharmacological manipulation showed that drug-induced sLFO-assessed arousal changes were accompanied by corresponding shifts in network connectivity and dynamics. This work identifies arousal as a prominent determinant of variability in functional brain networks, providing a new perspective on brain-based individual differences while offering an approach to measure arousal directly from fMRI data.

## INTRODUCTION

Disentangling individual differences in intrinsic brain function is crucial for advancing our understanding of the brain. These intrinsic functions lead to spontaneous fluctuations in measured brain activity. Despite the basic tenet that these spontaneous fluctuations reflect regionally specific neuronal activity and communication, predictive modeling has recently demonstrated that a large fraction of the spontaneous fluctuations in brain activity can be explained by physiological arousal alone^1^. While this work was carried out in rodents, a growing body of evidence using noninvasive brain imaging techniques, such as functional magnetic resonance imaging (fMRI), has begun to demonstrate a strong link between arousal and functional brain organization in humans as well^2–9^.

While a branch of fMRI research has directly mapped arousal-related brain networks or arousal-related brain fMRI signatures in humans^10,11^, more recent work has shown that physiological arousal is also closely intertwined with functional brain networks which were initially identified *independently* from systemic physiology. Specifically, evidence indicates that arousal is related to canonical functional brain networks, which are composed of spatially discrete brain regions that reliably co-activate to support specific cognitive, affective, or sensory functions and whose dysfunction is implicated across neuropsychiatric conditions^2,4,7,9,12–15^. For example, the Default Mode Network (DMN), involved in internally oriented attention, shows activation that precedes pupil dilation (a reliable marker of arousal) and this DMN activity in turn predicts suppression of the sensorimotor network (SMN)^15^. Others have shown that increased arousal is associated with DMN suppression and activation of a ‘Task Positive Network’^2^. Task Positive Networks include the SMN as well as the Salience Network (SN) and Dorsal Attention Network (DAN), which have been traditionally characterized as ‘task positive’ because they support externally oriented attention (and external attention was historically modeled during tasks). Collectively, these findings demonstrate that heightened arousal is related to stronger antagonism between functionally opposing (internal vs. external) networks, fitting with a body of literature that shows heightened arousal is related to stronger static anti-correlation between the DMN and externally oriented functional brain networks^5^. This establishes that heightened arousal strengthens the antagonistic balance between internal and external networks, and critically, it suggests that arousal may be fundamentally involved in functional network connectivity and dynamics. However, it is clear that this relationship is not fully understood, and a systematic investigation on the relationships between systemic physiological arousal and functional brain network organization is needed. Moreover, the field of functional imaging often treats these arousal-related processes as noise and disregards the primary role that these processes may have in functional brain network investigations.

A primary reason for disregarding arousal in fMRI neuroimaging studies is methodological constraints. It is challenging, and can be costly, to measure arousal independently of the fMRI scan. Further, even if arousal is measured, it is always measured somewhat by proxy, as defining “arousal” precisely has long been a challenge in the field. This is because arousal is a multidimensional construct encompassing both physiological and psychological components. Physiological arousal reflects changes in bodily systems such as heart rate, pupil diameter, respiration, and systemic blood flow, while psychological arousal captures subjective alertness or attentional engagement. These components often correlate but can diverge depending on context or individual differences. As such, measures of arousal often capture only a subset of the underlying processes and are susceptible to extraneous influences.

Here, we address both the challenge of measuring arousal in fMRI studies and the challenge of capturing a more central index of arousal by leveraging scientific advancements in our understanding of the distinct components of the BOLD signal. Specifically, we use the BOLD signal to concurrently measure systemic physiology in addition to the more conventional measurement of neuronal activation. One distinct component of the BOLD signal is the systemic low frequency oscillations (sLFO)^16^. The sLFO reflects slow fluctuations in blood oxygenation and volume, typically in the 0.009–0.15 Hz range, which can be measured in the brain and peripherally (for example, with pulse oximetry on a fingertip); these signals at distinct locations in the body are highly correlated, differing primarily in relative delay due to blood arrival times^16,17^. Importantly, the sLFO is physiologically distinct from task-evoked and resting state BOLD signals, driven by local neurovascular coupling, which reflect focal neuronal activity. Because of this, the sLFO has traditionally been treated as “physiological noise”. However, the sLFO itself carries meaningful information about autonomic function. Recent work demonstrates that the dynamic sLFO co-fluctuates with multiple indices of physiological arousal, including smooth muscle contractions, respiration, and pupil diameter^18^. This comprehensive investigation of the sLFO found that the sLFO is positioned to operate as a window into the function of the autonomic nervous system. Further, a sLFO derived metric—the mean sLFO amplitude—correlates strongly with several traditional indices of arousal (at R > 0.90 for each physiological index)^19^. These relationships collectively indicate that higher arousal is strongly associated with lower sLFO amplitude.

Notably, low frequency oscillations in BOLD fMRI (which includes the sLFO) have also been related to biological processes that may not, at first glance, appear related to arousal. For instance, the sLFO has been implicated in the movement and clearance of cerebral spinal fluid in humans^20–23^. However, this function likely arises from the same blood-borne, cardiopulmonary physiology that links the sLFO to arousal^22,24^. Thus, it is probable that the diverse physiological roles related to the sLFO reflect shared systemic mechanisms rather than distinct processes or evidence of a conflicting theory of sLFO function. Altogether, because the sLFO is simultaneously measurable within standard fMRI scans and centrally implicated in autonomic arousal, it provides a unique opportunity to investigate arousal’s relationship to functional brain networks at a large scale.

Equipped with this fMRI-derived arousal measure, we conducted a large-scale study to examine how sLFO-assessed arousal relates to both the spatial organization (network-network connectivity) and the time-varying activity of functional brain networks. Of note, because the mean sLFO amplitude has only been recently established as an index of arousal, we also replicated the following arousal x network investigation using alternative, more traditional indices of arousal. We first assessed relationships between the mean sLFO amplitude and network-network connectivity between all pairs of seven canonical brain networks. Given its predominant role at rest and known interactions with externally oriented networks, we then focused on the DMN. Based on prior work, we hypothesized that heightened arousal (lower sLFO) would be associated with stronger anti-correlations between the DMN and externally oriented networks. Next, we assessed how arousal is related to the temporal dynamics of individual brain networks. Time-varying network analyses capture fluctuations in network activity across the scan, providing insights into moment-to-moment brain states and their potential links to cognition and psychopathology^25–28^. By combining the sLFO index of arousal with dynamic network measures, our approach allows us to move beyond static connectivity and directly interrogate how arousal shapes both the spatial organization and the dynamic behavior of functional brain networks.

Finally, we examined these relationships following administration of methylphenidate and haloperidol, which are a catecholaminergic agonist and antagonist, respectively. This pharmacological manipulation allowed us to test whether the relationship between arousal and network properties was modulated by dopamine/norepinephrine signaling, two systems deeply involved in arousal^29–31^. By evaluating sLFO-network relationships across distinct catecholaminergic states, we can determine the extent to which these arousal effects are stable, neurochemically driven, and/or differentially expressed under heightened or reduced catecholaminergic tone. Taken together, this work positions physiological arousal as a key contributor to individual variability in brain network properties, offering a framework to integrate systemic physiology into large-scale neuroimaging research.

## RESULTS

### Results Overview

An overview of the present work is depicted in Figure 1. Please note that in all investigations on the sLFO as it relates to functional brain networks, we use only the mean sLFO amplitude as our sLFO measure of interest. This sLFO derivative specifically is an established index of arousal^19^. Because we only use this sLFO derivative in the following analysis, we often refer to mean sLFO amplitude as “mean sLFO” or “sLFO” for brevity.

**Figure 1.**
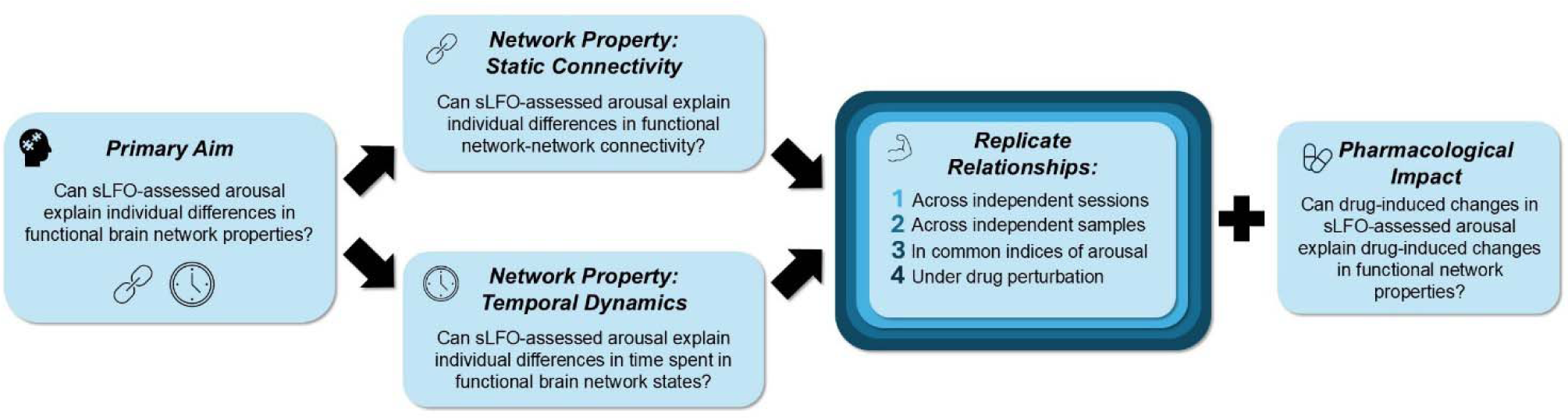
Overview of the present work. The primary aim of the present work is to understand whether arousal, measured as mean sLFO amplitude, may explain individual differences in functional brain network properties. Functional network properties investigated include both static network-network connectivity and network temporal dynamics. Relationships were replicated across multiple metrics.

### Mean sLFO x Functional Network-Network Connectivity

Analyses were conducted with data from the Human Connectome Young Adults Project (HCP-YA) which consists of two scan sessions of data per subject. We first confirmed that sLFO denoising reduced apparent global functional connectivity (Figure S1). After this confirmation, we began our analysis by testing whether the mean sLFO amplitude (i.e., the fraction of the LFO band variance in each voxel explained by the voxel wise sLFO regressor, averaged over the brain) was broadly related to functional network-network interactions. We tested 21 linear models relating the mean sLFO to Fisher r-to-z transformed network-network connectivity of each network-network interaction across 7 canonical brain networks, controlling for age, sex, phase encode direction, and scan session. Mean sLFO was significantly related to each network-network connectivity pair after Bonferroni correction for 21 comparisons (p<0.001; Table S1). R-values calculated from each of the 21 network-network connectivity x mean sLFO relationships are shown as 7x7 matrices in Figure 2A; across all network-network interactions, the sLFO was positively related to network-network connectivity.

**Figure 2.**
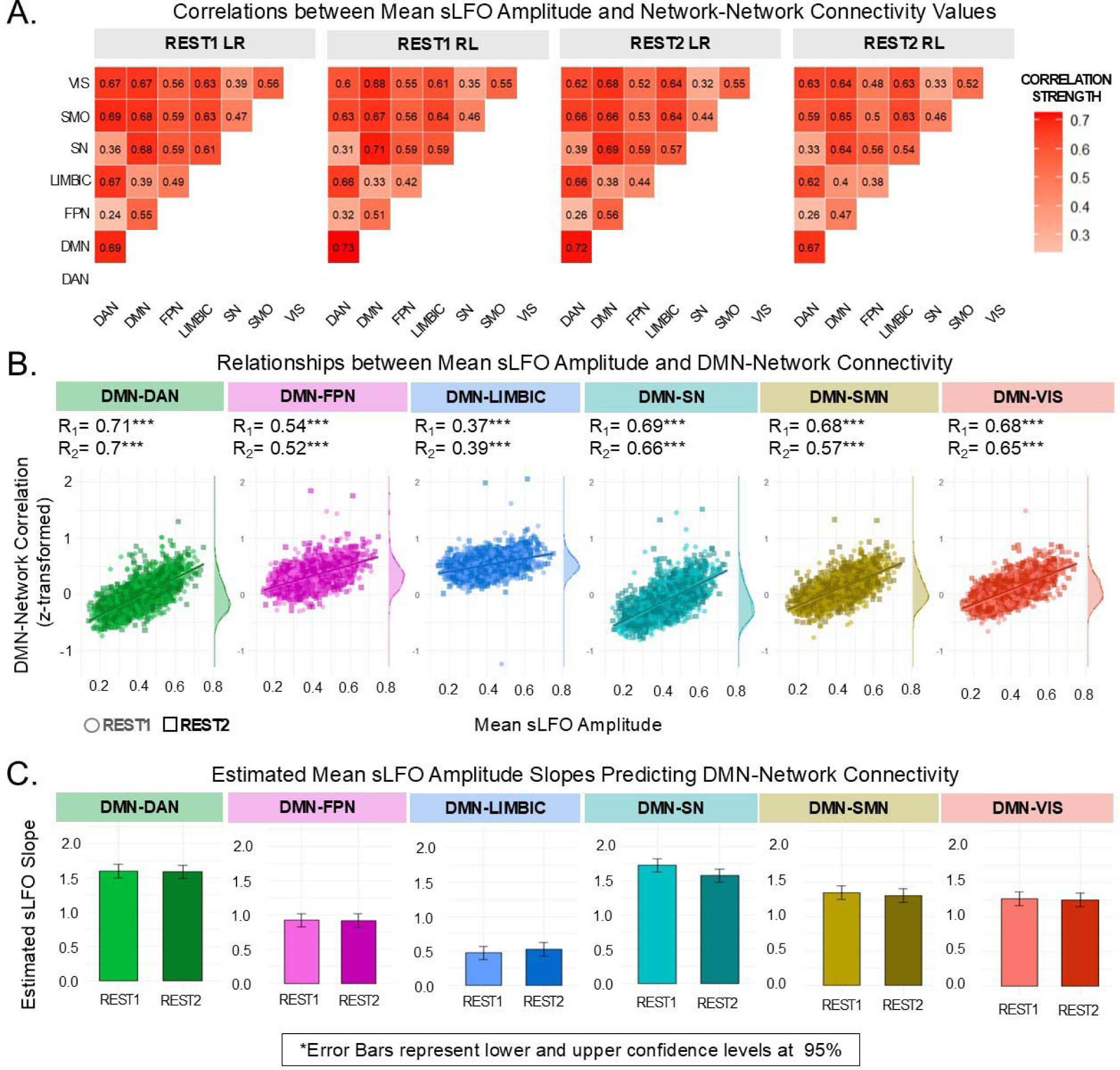
Mean sLFO amplitude explains a large proportion of individual differences in network-network connectivity. A. Correlations between mean sLFO amplitude and functional network-network connectivity values. B. Relationships between mean sLFO amplitude and DMN-network connectivity. Correlation value for session 1 is indicated as R_1_ and session 2 is indicated as R_2_. C. Estimated mean sLFO amplitude slopes predicting DMN-Network activity. Models controlled for age, sex, phase encode direction, and within subject random effects. DMN: Default Mode Network, DAN: Dorsal Attention Network, FPN: Frontoparietal Network, LIMBIC: Limbic network, SN: Salience Network, SMN: Somatomotor Network, VIS: Visual Network. *p*<0.001***

We then focused on how the DMN, the predominant functional brain network at rest, was communicating with other functional brain networks. We conducted a mixed linear model to test the relationship between mean sLFO and connectivity between the DMN and the following remaining six canonical brain networks: DMN-DAN, DMN-Frontoparietal Network (FPN), DMN-SN, DMN-Limbic Network, DMN-SMN, and DMN-Visual Network (VIS). Models controlled for age, sex, phase encode direction (AP vs. PA), and session (REST1 and REST2). There were main effects of mean sLFO (F_(1,8118.1)_=4711.57, p<0.001) and network (F_(5/2298.41)_=2487.43, p<0.001). There was also a significant mean sLFO x network interaction (F_(5/7857.16)_=164.72, p<0.001), indicating network-specific relationships between the sLFO and DMN-network connectivity. There were also main effects of sex (F_(1/460.93)_=4.27, p=0.04) and session (F_(1/8306.5)_=8.32, p=0.004) indicating an overall effect of these variables on network connectivity values. There was no main effect of phase encode direction (p>0.05). Of note, our session-based reproducibility analysis addressed whether the sLFO-DMN-network connectivity relationship changes as a function of session. Correlations between the mean sLFO and each DMN-network are shown in Figure 2B and model results are in Table S2.

All sLFO x DMN-network relationships exhibited the same directionality such that lower sLFO (indexing heightened arousal) was related to stronger DMN-network anti-correlation (Figure 2B). Thus, to investigate the sLFO x DMN-network interaction—namely, the variance in strength of sLFO x DMN-network relationships across the six networks—we conducted pairwise comparisons of the sLFO slopes predicting network connectivity across each DMN-Network (Figure 2C; Table S3). In session 1, this post-hoc analysis revealed that there was no significant difference in the sLFO slope predicting DMN-DAN correlation compared to the sLFO slope predicting DMN-SN correlation (z=2.032, p=0.32). There was also no significant difference in the sLFO slope predicting DMN-SMN connectivity compared to the sLFO slope predicting DMN-VIS connectivity (z=1.38, p=0.93). All other pair-wise comparisons of sLFO slopes significantly differed from each other when correcting for multiple comparisons (p<0.003), such that DMN-DAN and DMN-SN sLFO slopes were significantly greater than all other networks, and DMN-SMN and DMN-VIS sLFO slopes were significantly greater than DMN-FPN and DMN-Limbic sLFO slopes. These findings precisely replicated in session 2 data (DMN-DAN vs. DMN-SN z=0.3, p=0.99; DMN-SMN vs. DMN-VIS z=0.973, p=0.93); all other pairwise comparisons differed significantly when correcting for multiple comparisons (p<0.003; Table S3). Findings support the conclusion that the strength of relationships between 1) sLFO and DMN-DAN connectivity and 2) sLFO and DMN-SN connectivity do not differ from one-another and are significantly stronger than all other sLFO x DMN-network relationships.

### Reproducibility of Mean sLFO x Functional Network-Network Connectivity

To determine reproducibility 1) within the HCP-YA sample across separate scan days and 2) across independent samples of HCP-YA and an independent cohort of healthy adults (N=59), we conducted Bayes factor analysis. In contrast to testing a linear model for an sLFO x session or sample interaction, which is meant to test for significant *differences* (and thus is not suited to test or quantify *sameness*), Bayes factor analysis allows us to quantify the likelihood of sameness (i.e., reproducibility) or difference in the relationships between sLFO and brain network organization across sessions or samples.

Each Bayes factor was computed for each specific DMN-network (DMN-DAN, DMN-Limbic, etc.). For the first aim, testing reproducibility across sessions, we compared a null model (session, phase, age, sex, within subject covariates) relating network-network connectivity to mean sLFO to an alternate model which included a session x mean sLFO interaction. Based on interpretation from Lee and Wagenmakers^32^, Bayes factor analysis supported session reproducibility for all sLFO x DMN-network relationships at moderate-very strong levels (Table 1).

**Table 1.** Relationship between DMN-network connectivity and mean sLFO is reproducible in independent sessions and samples. Top: Bayes Factor analysis revealed that relationship between DMN-network connectivity and mean sLFO is reproducible across sessions. Bottom: Bayes Factor analysis reveals that relationship between DMN-Network connectivity and mean sLFO is reproducible in independent sample of healthy adults. DMN: Default Mode Network, DAN: Dorsal Attention Network, FPN: Frontoparietal Network, LIMBIC: Limbic network, SN: Salience Network, SMN: Somatomotor Network, VIS: Visual Network.

| **Between Sessions, within HCP-YA Sample Analysis** |  |
| --- | --- |
| **Brain Network State** | **Bayes Factor (H0/HA)** |
| DMN-DAN | 41.54 ± 9.69% |
| DMN-FPN | 35.3 ± 8.73% |
| DMN-LIMBIC | 6.62 ± 14.08% |
| DMN-SN | 3.8 ± 10.85% |
| DMN-SMN | 9.14 ± 8.63% |
| DMN-VIS | 38.92 ± 19.26% |
| **Between Samples Analysis (HCP-YA & N=59 Healthy Adults)** |  |
| **Between-Network Connectivity** | **Bayes Factor (H0/HA)** |
| DMN-DAN | 5.41 ± 10.32% |
| DMN-FPN | 5.845 ± 6.45% |
| DMN-LIMBIC | 5.18 ± 16.55% |
| DMN-SN | 9.07 ± 7.61% |
| DMN-SMN | 1.65 ± 7.99% |
| DMN-VIS | 1.85 ± 11.13% |

For the second aim, testing reproducibility between independent samples, we compared a complete null model (sample, age, sex, within subject covariates) relating network-network connectivity to mean sLFO to an alternate model which included a sample x mean sLFO interaction. Bayes factor analysis supported sample reproducibility for all sLFO x DMN-network relationships at anecdotal-moderate levels (Table 1).

### Mean sLFO x Functional Network Temporal Dynamics

We conducted a mixed linear model to test the relationship between mean sLFO and percent of time spent in brain network state across the eight brain network states previously derived via CAP analysis in this large sample of healthy young adults^33^. The brain network states, which align with known functional brain networks, are named as follows: frontoinsular-DMN (FI-DMN), FPN, DMN, DAN, SN-1, sensorimotor occipital (SMN), sensorimotor-DMN (SM-DMN), and SN-2 (visualization in Figure 3A).

**Figure 3.**
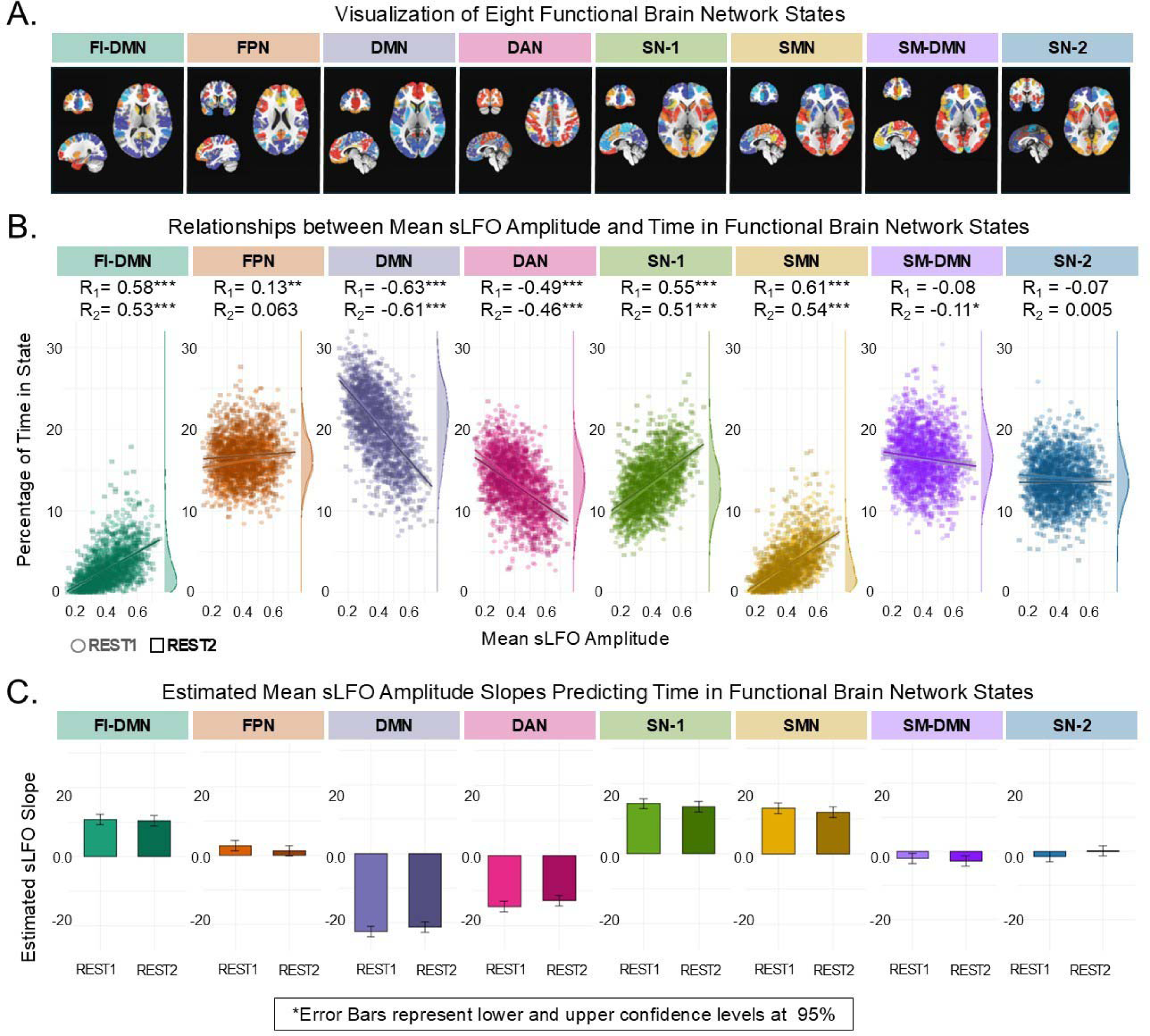
Mean sLFO amplitude explains a large proportion of individual differences in network temporal dynamics. A. Visualization of eight functional brain network states B. Relationships between mean sLFO amplitude and time in functional brain network states. Correlation value for session 1 is indicated as R_1_ and session 2 is indicated as R_2_. C. Slope of mean sLFO amplitude predicting time in functional brain network states. Models control for age, sex, phase encode direction and within subject random effects. FI-DMN: Frontoinsular-Default Mode Network State, FPN: Frontoparietal Network State, DMN: Default Mode Network State, DAN: Dorsal Attention Network State, SN-1: Salience Network State 1, SMN: Somatomotor Network State, SM-DMN: Somatomotor-Default Mode Network State, SN-2: Salience Network State 2. *p*<0.05*, *p*<0.01**, *p*<0.001***

There was a main effect of brain state (F_(7/6493.48)_=2,101.1, p<0.001) and a significant interaction between sLFO and brain state when relating to percentage of time in state (F_(7/14141.11)_=508.4, p<0.001). There were also significant brain state x sex (F_(8/3701.63)_=9.44, p<0.001), brain state x age (F_(8/3691.93)_=13.61, p<0.001), and brain state x session interactions (F_(8/11063.7)_=13.72, p<0.00). Correlations between the mean sLFO and time in each brain state are shown in Figure 3B; sLFO slope predicting time in each state is shown in Figure 3C, and each individual model result is in Table S4.

### Reproducibility of Mean sLFO x Functional Network Temporal Dynamics

Mirroring the approach used for testing reproducibility between mean sLFO and DMN-network connectivity, we used Bayes factor analysis to test for reproducibility in the relationships between sLFO and brain network temporal dynamics. In testing reproducibility between sessions, Bayes factor analysis revealed anecdotal-very strong evidence of between session reproducibility for all brain network states (Table 2). For testing reproducibility between samples, Bayes factor analysis supported sample reproducibility for the DMN, DAN, SM-DMN, and SN-2 at anecdotal-moderate levels while in contrast, there was very strong evidence of a sample effect for the FI-DMN and FPN, and there was weak anecdotal evidence of a sample effect for the SN-1 and SMN. This indicates brain state specificity in the reproducibility of the sLFO x brain state temporal dynamic relationships (Table 2).

**Table 2.**
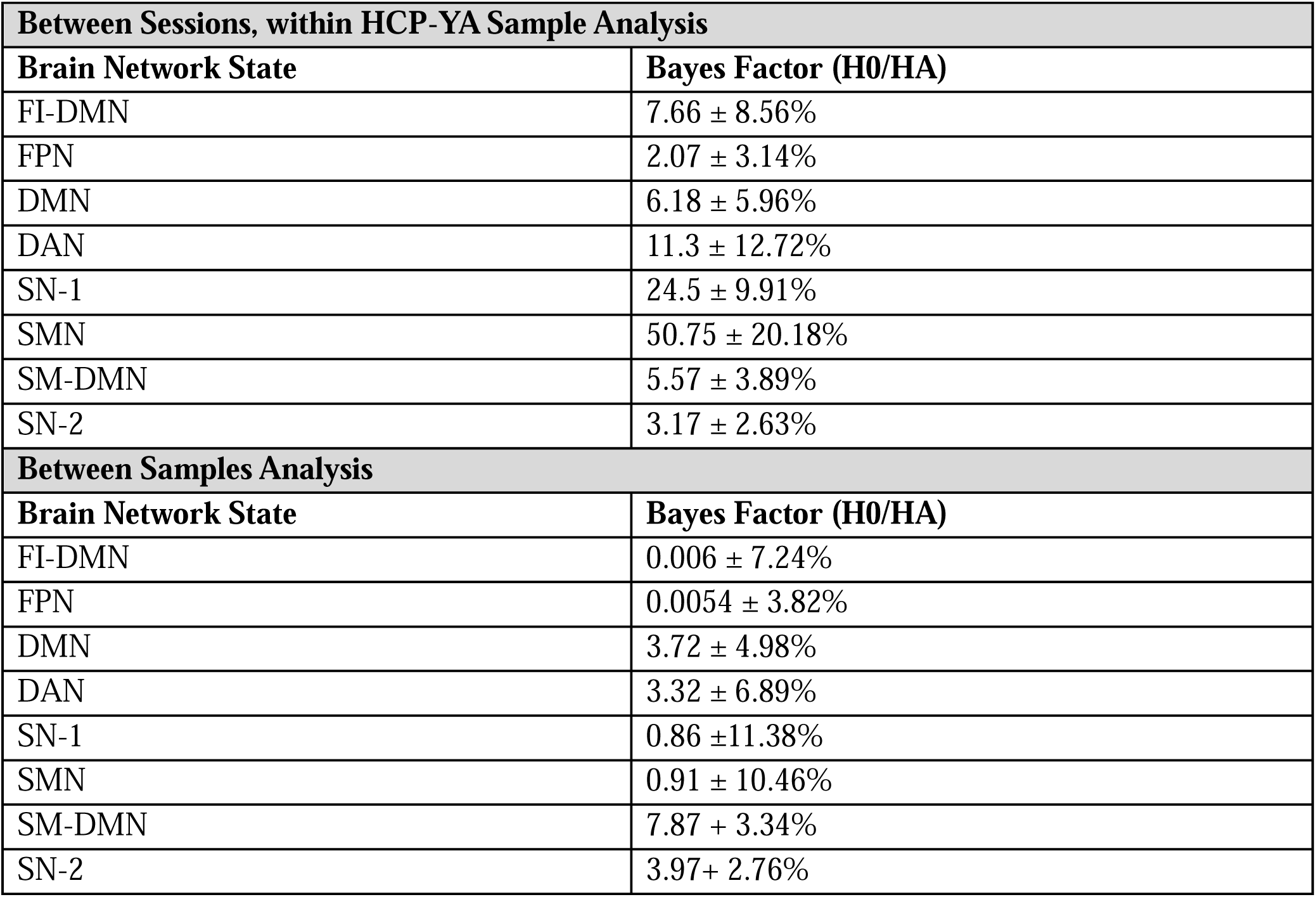
Relationship between functional network temporal dynamics and mean sLFO is reproducible in independent sessions and samples in a network-specific manner. Top: Bayes Factor analysis reveals that relationship between sLFO and FI-DMN, DMN, DAN, SN-1, SMN, SM-DMN, and SN-2 is reproducible across sessions. Bottom: Bayes Factor analysis reveals that relationship between network temporal dynamics and mean sLFO is reproducible in independent sample of healthy adults for DMN, DAN, SM-DMN and SN-2. For FI-DMN and FPN, Bayes Factors analysis reveals that a sample effect is supported. For SN-1 and SMN, a weak anecdotal sample effect is supported. FI-DMN: Frontoinsular Default Mode Network State, FPN: Frontoparietal Network State, DMN: Default Mode Network State, DAN: Dorsal Attention Network State, SN-1: Salience Network State 1, SMN: Somatomotor Network State, SM-DMN: Somatomotor-Default Mode Network State, SN-2: Salience Network State 2.

| **Between Sessions, within HCP-YA Sample Analysis** |  |
| --- | --- |
| **Brain Network State** | **Bayes Factor (H0/HA)** |
| FI-DMN | $7.66 \pm 8.56\%$ |
| FPN | $2.07 \pm 3.14\%$ |
| DMN | $6.18 \pm 5.96\%$ |
| DAN | $11.3 \pm 12.72\%$ |
| SN-1 | $24.5 \pm 9.91\%$ |
| SMN | $50.75 \pm 20.18\%$ |
| SM-DMN | $5.57 \pm 3.89\%$ |
| SN-2 | $3.17 \pm 2.63\%$ |
| **Between Samples Analysis** |  |
| **Brain Network State** | **Bayes Factor (H0/HA)** |
| FI-DMN | $0.006 \pm 7.24\%$ |
| FPN | $0.0054 \pm 3.82\%$ |
| DMN | $3.72 \pm 4.98\%$ |
| DAN | $3.32 \pm 6.89\%$ |
| SN-1 | $0.86 \pm 11.38\%$ |
| SMN | $0.91 \pm 10.46\%$ |
| SM-DMN | $7.87 \pm 3.34\%$ |
| SN-2 | $3.97 \pm 2.76\%$ |

A notable difference between our samples that may be at play in the lack of reproducibility is age range: 22-36 years in the original HCP-YA sample vs. 18-53 years in the independent sample. We thus conducted a post hoc analysis to test for an effect of age in the sLFO x brain state temporal dynamics relationships in the HCP-YA sample. We determined an effect of age for the FI-DMN, FPN, SN-1, and DAN states (see Supplementary Results; the effect is trend-level for FI-DMN and FPN). These brain states are three of the four states which exhibited evidence for a sample effect, lending to perhaps a biologically relevant effect of sample.

### Within Subject Changes in Mean sLFO and Functional Brain Network Properties

After demonstrating that individual variance in mean sLFO explains a large proportion of individual differences in brain network interactions and activity *between* subjects, we aimed to test whether intra-subject variance in mean sLFO explains variance in measures of brain function within subjects—in other words, we tested whether change in mean sLFO across the two scan sessions related to change in brain DMN-network connectivity and network temporal dynamics across the two scan sessions.

For DMN-network connectivity, there was a significant effect of ΔsLFO (F_(1/4539.56)_=1947.68, p<0.001) and a significant ΔsLFO x network interaction (F_(5/5070.35)_=112.64, p<0.001) in relating to ΔDMN-network connectivity (Figure 4A). There was also a significant effect of phase encoding direction (F_(1/5315.71)_=132.41, p<0.001). However, in an additional model, we confirmed there was no ΔsLFO x phase encode direction interaction, indicating that the relationship between change in sLFO and change in network connectivity is consistent across phase encode directions (p>0.05). Mirroring between subject findings, each ΔDMN-network relationship was positively related to ΔsLFO (Figure 4A). The strength of correlations between ΔsLFO and ΔDMN-network connectivity varied by network in a manner which also precisely aligned with the between subject correlations (Figure S2; Table S5), such that ΔsLFO slopes predicting ΔDMN-DAN and ΔDMN-SN did not differ (z = 0.749, p=0.97) and were significantly stronger relative to other slopes (p<0.003; Figure S2; Table S4). The ΔsLFO slopes predicting ΔDMN-SMN and ΔDMN-VIS also did not significantly differ (z=0.346, p=0.99) and were significantly stronger relative to ΔsLFO slopes predicting ΔDMN-FPN and ΔDMN-Limbic.

**Figure 4.**
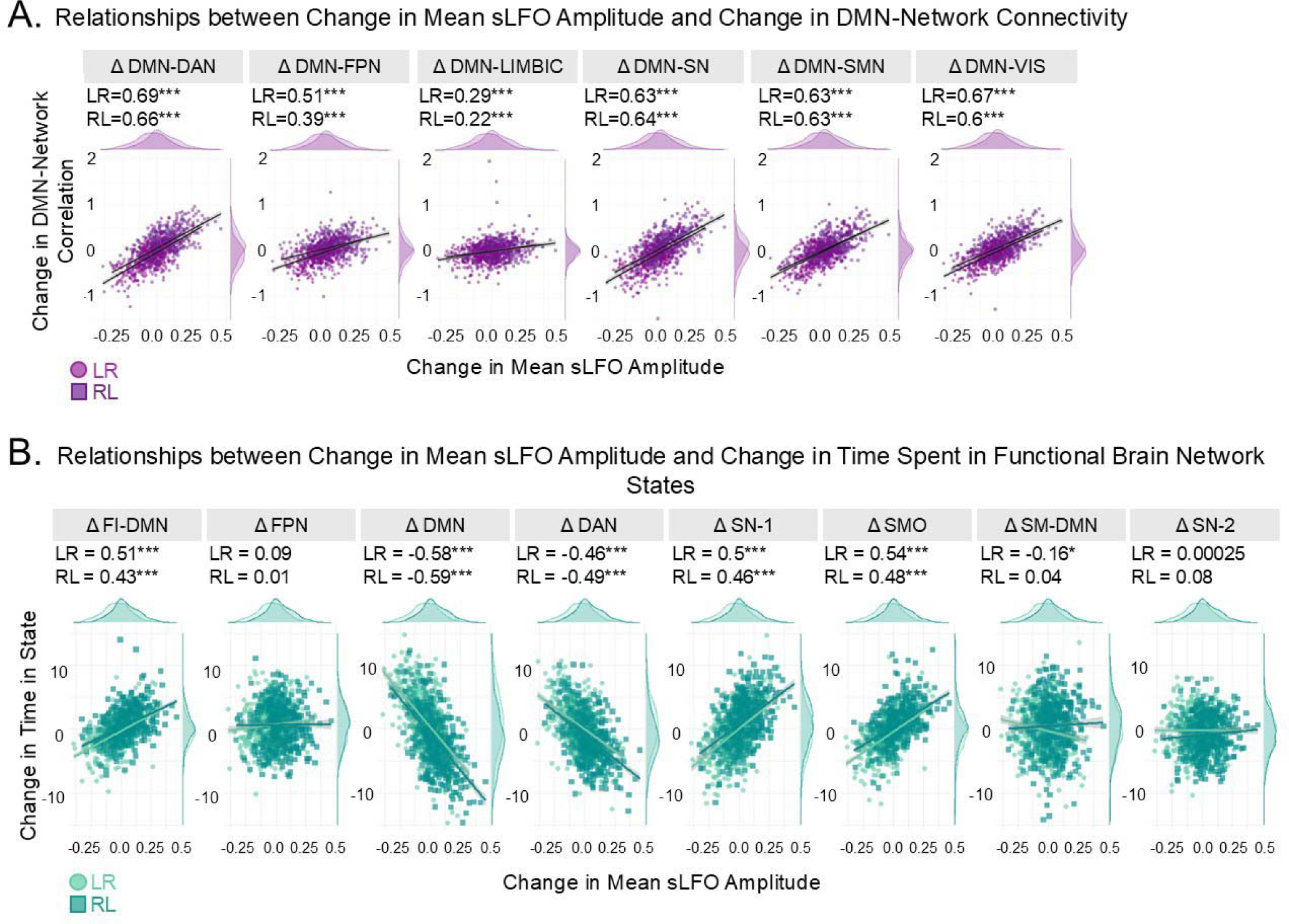
Intra-subject variability in mean sLFO amplitude explains a large proportion of intra-subject variability in network-network connectivity and network temporal dynamics. A. Relationship between change in mean sLFO amplitude and change in DMN-network connectivity. B. Relationship between change in mean sLFO amplitude and change in brain network temporal dynamics. For both A and B, change was measured within subject from scan session 1 to scan session 2 for two phase encodings (LR/RL); corresponding R-values are noted. DMN: Default Mode Network, DAN: Dorsal Attention Network, FPN: Frontoparietal Network, LIMBIC: Limbic network, SN: Salience Network, SMN: Somatomotor Network, VIS: Visual Network. FI-DMN: Frontoinsular Default Mode Network State, SM-DMN: Somatomotor-Default Mode Network State. *p*<0.05*, *p*<0.01**, *p*<0.001***

Similarly, the relationships between change in sLFO and change in network temporal dynamics across scan days largely mirrored the between subject findings (Figure 4B; Table S6). In the main model relating to Δtime in state, there was a significant effect of brain state (F_(7/14744)_=6.97, p<0.001) and there was a significant ΔsLFO x brain state interaction (F_(7/14744)_=460.58, p<0.001). There were also brain state x sex (F_(8/14744)_=3.02, p=0.002) and brain state x phase encode direction interactions (F_(8/14744.00)_=25.09, p<0.001). Post-hoc analysis revealed that ΔsLFO was related to Δtime spent in the FI-DMN, DMN, DAN, SN-1, and SMN (p<0.001). Change in mean sLFO was not related to change in time spent in the FPN or SN-2 state (p>0.05). In contrast to the between subject findings, change in mean sLFO was weakly related to time spent in the SM-DMN (p_corr_=0.025), however this was not consistent across runs (LR phase R = -0.16, p<0.001; RL phase R = 0.038 p>0.05).

### Confirming Relationships between Functional Brain Networks and Physiology

Having determined robust relationships between the mean sLFO—our index of systemic physiological arousal—and functional brain network properties, we sought to confirm that the relationships between mean sLFO and 1) DMN-network connectivity and 2) network temporal dynamics aligned with physiology x brain relationships using more traditional indices of arousal available in our dataset: respiration volume per unit time (RVT) and percent of heart rate variability (HRV) in high frequency range (HRV-HF) and low frequency range (HRV-LF).

There was a main effect of RVT on DMN-network connectivity (F_(1/7915.26)_=136.24, p<0.001) and a significant RVT x network interaction (F_(5/7360.72)_=9.4, p <0.001). There was a main effect of HRV-LF on DMN-network connectivity (F_(1/7840.69)_=182.56, p<0.001) and an HRV-LF x DMN-network interaction (F_(5/6352.69)_=7.6, p<0.001). There was also a main effect of HRV-HF on DMN-network connectivity (F_(1/7513.89)_=23.31, p<0.001) however there was no HRV-HF x network interaction indicating a non-network specific relationship between HF HRV and DMN-network connectivity (p>0.05). The directionality of the relationships are as follows: RVT and HRV-HF are negatively related to DMN-network connectivity; HRV-LF is positively related to DMN-network connectivity. Thus, these findings confirm that, as measured by the mean sLFO and as consistent with the literature^5^, increased arousal (now indexed as higher RVT and HRV-HF and lower HRV-LF) is related to stronger DMN-network anti-correlation. Post-hoc analysis also confirmed that significant network-specific differences in the strength of the alternate indices of arousal predicting DMN-network connectivity which were tested/detected with these indices aligned with the network-specific differences in the strength of the sLFO slopes predicting DMN-network connectivity (Figure S3A; Table S7-S8).

In testing relationships between alternate measures of arousal and brain network temporal dynamics, the following interactions were detected in three separate models: brain state x RVT (F_(7/13456.6)_=23.76, p<0.001), brain state x HRV-LF (F_(7/13196.18)_ =32.64, p<0.001), brain state x HRV-HF (F_(7/12246.98)_ =4.88, p<0.001). Network-specific relationships between alternate indices of arousal and temporal dynamics again aligned with sLFO relationships in both strength and directionality (Figure S3B; Table S9).

### Mean sLFO, Functional Brain Networks, and Catecholaminergic Pharmacology

After determining relationships between the mean sLFO and functional brain network properties in sample of healthy adults under homeostatic conditions, we inquired whether such relationships would remain stable in healthy adults under pharmacological perturbation. This sample underwent three fMRI scan sessions, on separate days, under the following drug conditions: placebo, methylphenidate, and haloperidol (in a double-blind, randomized fashion). This pharmacological manipulation allowed us to probe mean sLFO x brain network relationships under drug conditions with antagonistic effects on both arousal and catecholamines: methylphenidate is a psychostimulant and a dopamine and norepinephrine agonist (through blocking reuptake of dopamine and norepinephrine), and haloperidol has sedative properties and is a dopaminergic D2 receptor antagonist.

In a linear model which controlled for age and sex, there were main effects of sLFO (F_(1/668.94)_=119.45, p<0.001), network (F_(5/321.60)_=173.07, p<0.001), drug (F_(2/695.48)_=4.17, p=0.016), and age (F_(1/56.17)_=5.65, p=0.021) on DMN-network connectivity. There were no significant sLFO x drug (p>0.05) or network x drug (p>0.05) interactions. There was no significant drug x sLFO x network interaction, indicating that drug does not significantly alter the relationship between mean sLFO and DMN-network connectivity (p>0.05; Tables S10-S11).

For network temporal dynamics, in a linear model which controlled for age and sex, there was a main effect of brain state (F_(7/479.76)_=296.73, p<0.001) and there were interactions of sLFO x brain state (F_(7/1235.96)_=28.88, p < 0.001), drug x brain state (F_(14/970.60)_=3.74, p<0.001) and age x brain state (F_(8/443.90)_=1.97, p=0.049) on percent time in state. Again, there was no three-way sLFO x brain state x drug interaction, indicating that the mean sLFO x brain network temporal dynamics relationship remains consistent across pharmacological perturbation (F_(14/1174.73)_=1.48, p>0.05; Table S12-S13).

Having determined that the relationships between sLFO and brain network organization are not significantly modulated by drug, we set out to test whether drug-induced changes in sLFO were related to drug-induced changes in functional network properties. For this analysis, we homed in on the DMN, DAN, and SN. Our focus on DMN, DAN, and SN was guided by both methodological and scientific rationales. From a methodological standpoint, our HCP-YA sample showed that relationships between the sLFO and DMN, DAN, SN were often the strongest relative to other networks and were largely reproducible, including existing in the pharmacological manipulation sample (Figure S4). From a scientific perspective, there is robust evidence that the DMN, DAN, and SN (and their interactions) hold catecholamine-related functional and clinical relevance^34–41^. As such, we were interested in how change in 1) DMN-DAN and DMN-SN connectivity and 2) DMN, DAN, and SN temporal dynamics under catecholaminergic drug compared to placebo relate to change in mean sLFO under catecholaminergic drug compared to placebo.

In a simple correlation test, we found that drug-induced changes in sLFO were related to drug-induced changes in DMN-DAN connectivity (methylphenidate R=0.49, p<0.001; haloperidol R=0.41, p<0.001) and were related to drug-induced changes in DMN-SN connectivity (methylphenidate R=0.51 p<0.001; haloperidol R=0.52 p<0.001) (Figure 5A). The positive direction of these relationships indicates that drug-induced decrease in sLFO (i.e. an increase in arousal) is related to drug-induced strengthening of network-network anti-correlation, whereas drug-induced increase in sLFO (decrease in arousal) is related to drug-induced weakening of network-network anti-correlation. We also demonstrated that drug-induced changes in mean sLFO is related to drug-induced changes in the temporal dynamics of DMN, DAN, and SN—specifically change in the time-varying dominant activation of these networks. Drug-induced decrease in sLFO (increase in arousal) was related to spending more time in DMN (methylphenidate R = -0.45, p<0.001; haloperidol R = -0.54, p<0.001) and DAN (methylphenidate R = - 0.46, p<0.01; haloperidol R = -0.45, p<0.001) and was related to spending less time in SN (methylphenidate R = 0.33 p<0.05; haloperidol R = 0.39 p<0.01) (Figure 5B).

**Figure 5.**
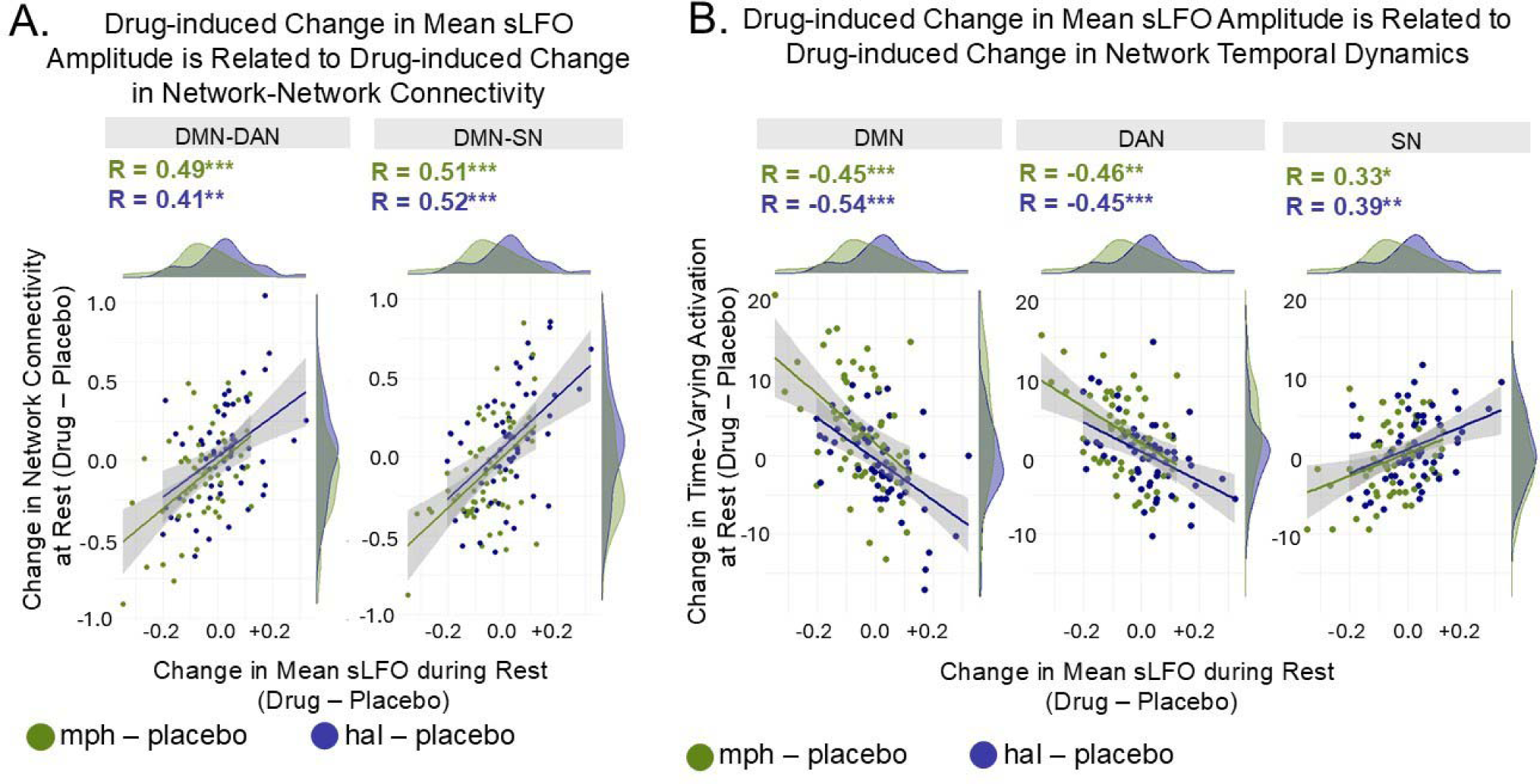
Drug-induced change in mean sLFO amplitude explains a large proportion of variability in drug-induced changes in functional network properties. A. Drug-induced change in mean sLFO during rest is related to drug-induced impact on DMN-DAN connectivity and DMN-SN connectivity. The positive direction of the relationship indicates that drug-induced decrease in sLFO (increase in arousal) is related to strengthening of DMN-DAN and DMN-SN anti-correlation, whereas drug-induced increase in sLFO (decrease in arousal) is related to weakening of DMN-DAN and DMN-SN anti-correlation. B. Drug-induced change in mean sLFO is related to drug-induced change in time spent in the DMN, DAN, and SN. DMN: Default Mode Network, DAN: Dorsal Attention Network, SN: Salience Network. *p*<0.05*, *p*<0.01**, *p*<0.001***

## DISCUSSION

Our findings demonstrate that large-scale functional brain network properties are strongly shaped by individual differences in systemic physiological arousal. By separating the systemic low-frequency oscillation (sLFO) from focal neuronal activity within the BOLD signal, we show that mean sLFO amplitude, a robust fMRI-derived index of arousal, explains a substantial proportion of variance in both functional network-to-network connectivity and time-varying network dynamics. Experimentally induced changes in arousal via pharmacological manipulation correspond with expected alterations in high-order network properties, while sLFO-network relationships remain consistent across sessions, samples, and pharmacological states. Moreover, relationships between arousal and brain networks measured via more traditional indices mirror the sLFO-network relationships found here. Together, these findings link individual differences in arousal to intrinsic brain function, highlighting the importance of accounting for arousal when interpreting spontaneous fMRI signals in studies of cognition, behavior, and psychopathology.

Previous work has established that the mean sLFO amplitude is tightly and inversely related to physiological arousal^18,19^. Extending this literature, we find that that lower sLFO (higher arousal) is consistently related to heightened anti-correlation between the DMN and task-positive networks such as the DAN and SN, which is consistent with prior studies linking pupil diameter and EEG indices of arousal to greater network-network anti-correlation^5,8^. Further, using more traditional physiological indices such as heart rate variability and respiration volume, we show that in our sample, traditional measures of arousal replicate the arousal-network relationships identified via mean sLFO amplitude. This provides convergent evidence that the mean sLFO amplitude captures biologically meaningful fluctuations in arousal. Importantly, all these relationships exist *after* regressing out the temporal sLFO signal from the BOLD data used to define network properties. This is critical as it removes the concern that these signals themselves might drive apparent links between arousal and network connectivity and dynamics.

Our analyses further demonstrate that interactions between higher-order cognitive networks, particularly the DMN-DAN and DMN-SN connectivity, show the *strongest* relationships with physiological arousal, which extends the work of others indicating that systemic physiology has a strong association with brain regions supporting cognitive function^4^. These cognitive networks are typically functionally opposing: the DMN is engaged during internally oriented attention, self-referential thought, and mind-wandering, whereas the DAN and SN are preferentially active during externally oriented attention and goal-directed task engagement^42–44^. The observed link between DMN–DAN / DMN–SN interactions and arousal suggests that heightened arousal, as indicated by a lower mean sLFO amplitude, may bias the brain toward different attentional modes—reflecting a functional segregation between internally-oriented (DMN) and externally-oriented (DAN, SN) processes, a principle that is consistent with prior models of network antagonism^45,46^. Given that imbalances among these same networks are central to widely used models of psychopathology (e.g., Menon’s triple network model^47^), our findings suggest that arousal measured via the sLFO may be an important physiological factor influencing intrinsic brain-state variation, which could have implications for cognitive performance and vulnerability to mental disorders.

Beyond static network connectivity, we show that mean sLFO amplitude strongly relates to time-varying brain network dynamics. Specifically, higher arousal corresponds to increased time spent in the DMN and DAN, which is a metric associated with heightened anti-correlation between these networks^36^. This finding supports classical models linking arousal to increased attentional state^48^ and builds on these models by revealing a novel association between arousal and the dynamics of attentional brain networks.

Interestingly, heightened arousal was associated with reduced time spent in the SN. This may reflect the SN’s role in coordinating transitions between attentional states: when arousal is high, attentional demands may be more effectively met due to increased time in the DMN and DAN, reducing the need for SN-mediated monitoring^7,9,49,50^. He et al. (2023) also found that heightened arousal was linked with increased switching from the SN into brain networks, including the DAN. Taken together, these findings support the possibility that, under heightened arousal, the SN primarily functions to initiate engagement of attentional networks, resulting in greater occupancy of these states and less time in the SN.

We go on to demonstrate that sLFO-network relationships are stable under pharmacological perturbation and that drug-induced changes in sLFO are strongly related to drug-induced changes in network properties. The positive directionality of these findings fits with known drug effects: for methylphenidate, a psychostimulant and attention enhancing medication, drug-induced decrease in sLFO (increase in arousal), is directly related to the strengthening of DMN-DAN and DMN-SN anti-correlation and the increase in time spent in attentional states. For haloperidol, a D2 antagonist with sedative properties and impairing effects on cognition, drug-induced increase in sLFO (reduction in arousal) is directly related to weakened anti-correlation. These results highlight that individual variability in drug-induced changes to cognitive network interactions is largely explained by changes in systemic physiological arousal, emphasizing the potential of the sLFO as a mechanistically grounded marker for predicting brain-based responses to psychoactive drugs and pointing to the largely unexamined role that systemic physiology may play in neurobiological responses to drugs.

The effect sizes observed here are striking: for the DMN results, individual differences in DMN-DAN connectivity correlates with the sLFO at R = 0.71, and time spent in the DMN correlates with the sLFO at R=0.63. Relationships of this strength suggest that mean sLFO amplitude may serve as a scalable, non-invasive, biologically informative proxy for network-level brain function and connectivity^10^. This high explanatory power likely stems from the sLFO’s central role as a core autonomic-regulated signal, rather than a system-specific proxy vulnerable to local confounds. Theoretical and empirical work supports the sLFO as a central correlate of vascular tone and autonomic function, coupling with smooth muscle contractions, respiration, and other arousal-related processes^18^. Moreover, prior work demonstrates that the mean sLFO amplitude is tightly associated with multiple traditional indices of arousal such as pupil diameter, HRV, and respiration, measured from peripheral locations^19^, highlighting its robustness and systemic relevance. Indeed, we observed that the sLFO x brain network relationships are less variable compared to alternate indices of arousal. Taken together, the sLFO may capture arousal-state more precisely compared to arousal indices that can vary by individual differences in specific systems, allowing for high explanatory power.

Although we demonstrate robust relationships between the sLFO and network properties in healthy adults, it remains to be seen whether the sLFO and network properties remain tightly coupled across age, neuropsychiatric conditions, or other perturbations to homeostatic brain physiology. Indeed, our secondary findings point to both stability and sensitivity of the sLFO-network relationship: we demonstrated that acute pharmacological manipulation does not alter sLFO-network coupling and instead shifts the sLFO and network properties in tandem, lending to the stability of sLFO-network relationships. However, we also showed preliminary evidence of an age by sLFO interaction in relating to network properties, such that increasing age is related to weakening of the sLFO-network relationship. Taken together, these findings suggest that the coupling between sLFO and network properties may be stable or disrupted depending on specific biological or clinical conditions. Given the strong relationship between the sLFO and brain networks established here in healthy young adults, determining how specific conditions may modulate the sLFO-network relationship holds potential to advance our understanding of brain-body coupling across the age span and disease.

Further, the ability to measure sLFO peripherally via pulse oximetry^17^, raises the intriguing possibility of developing biomarkers of cognitive network function that are non-invasive and feasible to assess for routine clinical use. To this aim, future work would need to validate whether peripheral measures of sLFO align with cognitive network properties. This confers considerable research effort to move towards such a biomarker—however, the potential of an affordable, easy-to-use proxy of brain function is nonetheless compelling.

A limitation of the work here is that not all sLFO x brain state temporal dynamics relationships were reproducible. While four of the eight states showed evidence of reproducibility, the remaining four supported evidence for a sample effect. A notable difference between our samples was age range: 22-36 years in the original sample vs. 18-53 years in the independent sample. With this in mind, we conducted a post hoc analysis to test for an effect of age in the sLFO x brain state temporal dynamics relationships. We determined an effect of age for the FI-DMN, FPN, SN-1, and DAN states which are three of the four brain states which exhibited evidence for a sample effect. Thus, it may be that the sample effect reflects a biologically relevant shift in the sLFO x brain network temporal dynamics relationship across age, particularly as there are known effects of age on neurovascular function^51^, functional brain network properties^52^, and a demonstrated effect of arousal on age-related changes in functional networks^53^. This invokes a second limitation: that the present sample did not have an age range large enough to fully understand how age may modulate sLFO x network relationships. Future work using datasets such as HCP-aging or the Adolescent Brain and Cognitive Development (ABCD) study to understand the effect of age on the sLFO-brain network relationships is of interest. However, we provide the first evidence of robust sLFO x brain networks relationships, creating a foundation on which to build to understand how changes in brain function, such as those which occur across age or in disease, impact the sLFO-network relationships.

In conclusion, our results demonstrate that systemic physiological arousal, as indexed by mean sLFO amplitude, is a primary determinant of individual differences in both static and dynamic functional network organization. This work establishes a framework for integrating systemic physiology into large-scale neuroimaging studies, reveals mechanistic links between arousal and cognitive brain function, and opens new avenues for precision medicine approaches that leverage intrinsic physiological signals to predict individual brain function and responses to pharmacological interventions.

## METHODS AND MATERIALS

### Samples

*Human Connectome Project Young Adults* (HCP-YA) Cohort included participants from the HCP 1200 subject release. Data was used from 462 participants from the HCP-YA sample who were previously assessed given they have no current or family history of certain psychiatric illness or substance use (181 male, 281 female; age 28.66 <u>±</u> 3.65 years)^33^. Resting-state scans were captured on two independent days, with two scans per day recorded using different phase encode directions: right to left (RL) and left to right (LR). All participants provided written consent prior to participation.

*Acute Drug Administration Cohort* included 59 healthy individuals (17 male, 42 female, aged 18-55). Participants with mental and physical health diagnoses and/or medications that interfere with the BOLD signal or alter metabolism of catecholaminergic agents were excluded. Participants were scanned in a resting-state paradigm following administration of 20 mg methylphenidate, 2 mg haloperidol or placebo–timed in accordance with drug peak–using a randomized, counterbalanced, double blind design. The study was reviewed and approved by the institutional review board of the National Institutes of Health and all participants provided written informed consent prior to participation.

### fMRI Acquisition

*HCP-YA Resting-State Scan:* Details of HCP-YA scan and opensource preprocessed data can be found elsewhere^54,55^. Briefly, data were collected using a Siemens 3T Skyra scanner with a 32-channel head coil. Scan parameters were as follows: TR = 0.72 s, TE = 33.1 ms, flip angle = 52°, field of view = 280 x 180 mm^2^, matrix = 140 x 90, echo spacing = 0.58 ms, bandwidth = 2,290 Hz px^-1^. Slice thickness was set to 2.0 mm, 72 slices and 2.0 mm isotropic voxels, multiband acceleration factor = 8. Total scan time was 14.4 minutes for each resting state scan—one each visit. Each run was completed with oblique axial acquisitions and alternated between phase encoding left-to-right (LR) and phase encoding right-to-left (RL). The opensource HCP minimal preprocessed resting-state fMRI data was used in this work^54^. RIPTiDe was conducted on the cleaned data to quantify and remove sLFO signal driven by global blood-based physiology^56,57^. Note that negative correlations are an unavoidable consequence of global signal regression, but this is not the case for sLFO regression via RIPTiDe. This is because RIPTiDe adjusts for the time delay of the sLFO signal, allowing for full removal of the sLFO signal specifically and leaving little to no inverted signal in the residuals^58^ (e.g., spurious anti-correlations).

*Pharmacological Manipulation Cohort Resting-State Scan:* Data were collected using a Siemens Trio 3T scanner with a 12 channel RF coil. The 8-min resting state scan was acquired with gradient-echo, echo planar images using oblique axial scans 30° from AC-PC with AP phase encoding and the following parameters (TR = 2 s, TE = 27 ms, flip angle = 78°, voxel resolution = 3.4375 × 3.4375 × 4 mm). Resting-state fMRI data was processed using FMRIB software library (FSL6.0.0). Images underwent brain extraction, registration, spatial smoothing (6 mm), high-pass temporal filtering (100 s), and motion correction via MCFLIRT. Following motion correction, Independent Component Analysis via FIX was employed using a training set derived from the current data to further clean data^59,60^. Next, RIPTiDe was conducted on the data to quantify and remove temporal sLFO signal driven by global blood-based physiology. For functional connectivity analysis, TRs with motion greater than 0.35mm (Euclidean distance) were discarded.

### Deriving the Mean sLFO Amplitude

We derived the mean brain-wide sLFO amplitude first by running regressor interpolation at progressive time delays (RIPTiDe^56,57^) using the open source “rapidtide” suite (https://github.com/bbfrederick/rapidtide doi:10.5281/zenodo.17633117). RIPTiDe was conducted to derive mean sLFO amplitude prior to FIX correction as such correction can interfere with sLFO extraction. RIPTiDe outputs a brain-wide map of squared maximum correlation for each brain voxel, where the max correlation is the correlation between the ‘probe’ signal (global signal) and each voxel’s BOLD signal at the optimal time delay (i.e., at the time delay which allows for maximum correlation). The squared correlation is the fraction of the variance in the voxel explained by the sLFO signal. The mean brain-wide sLFO amplitude is defined as the average of the brain-wide max correlation squared. All subsequent analysis on sLFO x functional brain network relationships <u>only</u> use the mean brain-wide sLFO amplitude measure of the sLFO, as this sLFO derivative specifically is the established index of arousal^19^.

### Functional Network-Network Connectivity

To derive network-network functional connectivity values, we extracted mean time courses of each of the seven functional brain networks using the Yeo 7-network parcellation atlas^61^ and FSL (fslmeants) or AFNI (3dMean). We derived the Fisher r-to-z transformed correlation values between each pair of network-network time courses (21 correlations per subject). For HCP-YA, r-to-z correlation values were derived for each scan session and each phase encode direction (four total). For the pharmacological manipulation sample, r-to-z correlation values were derived under each drug condition.

### Mean sLFO x Functional Network-Network Connectivity

To test for relationships between the mean sLFO and network-network functional connectivity broadly, we began by conducting 21 linear mixed models which examined the relationship between sLFO and network-network connectivity while controlling for age, sex, scan session, phase, and within subject random effects, using the HCP-YA sample.

We then focused on the interactions between the DMN and other functional brain networks, given the importance of DMN-network connectivity in psychopathology, cognition, and demonstrated relationships between DMN-network connectivity and physiological arousal^2,5,8,39,41,47,62^. We conducted a linear mixed model which tested the interaction between sLFO x DMN-network (DMN-DAN, DMN-FPN, DMN-Limbic, DMN-SN, DMN-SMN, DMN-VIS) in relating to functional connectivity, while controlling for main effects of sLFO, DMN-network, age, sex, scan session, phase encode direction, and within subject random effects. This was carried out using the mixed function of the afex package in R. After finding a significant sLFO x DMN-network interaction, we conducted post-hoc analysis to compare the strength of sLFO slopes predicting DMN-network functional connectivity across the six DMN-network relationships, resulting in 15 pairwise-tests. This post-hoc analysis was conducted using the emtrends function of the emmeans package in R; significance was corrected for multiple comparisons using Bonferroni method.

### Reproducibility of Mean sLFO x Functional Network-Network Connectivity

To confirm reproducibility of relationships between mean sLFO and DMN-network correlations across separate scan days within the HCP-YA sample, we used Bayes factor analysis. For each DMN-network (DMN-DAN, DMN-Limbic, etc), we compared a null model to an alternate model. The null modeled the relationship between mean sLFO and network-network connectivity while controlling for session, age, sex, and within subject random effects. The alternate model solely differed from the null by including a mean sLFO x session interaction. Bayes factors which favor the null (BF>1) provide evidence for reproducibility as the sLFO slope would not vary by session. In contrast, favoring the alternate (BF<1) provides evidence for a session-based difference in the relationship between sLFO and DMN-network connectivity. Similarly, we used Bayes factor analysis to determine reproducibility of sLFO x DMN-network relationships across independent samples: the HCP-YA sample and our sample of 59 healthy adults under placebo (from the pharmacological manipulation trial). In this case, the null modeled the relationship between mean sLFO and DMN-network connectivity while controlling for sample, age, sex, and within subject random effects. The alternate model solely differed from the null by including a sLFO x sample interaction. Thus, here a Bayes factor which supports the null provides evidence for reproducibility across samples. In contrast, a Bayes factor which supports the alternate provides evidence of a sample-based difference in the relationship between sLFO and network-network connectivity. To conduct Bayes Factor analysis, we used the BayesFactor package in R and used Monte Carlo integration with the number of sampling iterations equal to 100,000. Bayes factors were qualitatively interpreted based on Lee and Wagenmakers^32^, with the following evidence thresholds: anecdotal (1-3), moderate (3-10), strong (10-30), and very strong (30-100). Reciprocal values indicate evidence of equal strength in the alternative direction.

### Functional Network Temporal Dynamics

To derive time-varying activity of functional brain networks, we conducted co-activation pattern (CAP) analysis. Specifically, we applied 8 CAPs which were previously defined by the HCP-YA sample, have been shown to be reliably active at rest, and align with known functional brain networks, including the DMN, DAN, FPN, SN, and SMN^33^. Given the spatial overlap between these data-driven CAPs and functional brain networks, we refer to the CAPs as “functional brain network states”. In conducting CAP analysis on resting-state data, we derived percentage of time spent in each brain network state at rest as our metric of interest.

### Mean sLFO x Functional Network Temporal Dynamics

To test relationships between the mean sLFO and percent of time in brain network state, we conducted a linear mixed model which tested for an interaction between sLFO and brain state (FI-DMN, FPN, DMN, DAN, SN-1, SMN, SM-DMN, SN-2) in relating to percentage of time in brain network state, while controlling for age, sex, scan session, phase encode direction, and within subject random effects. This was carried out using the mixed function of the afex package in R. Finding a significant sLFO x brain state interaction, we ran a linear model testing relationship between sLFO and percentage of time in state for each brain state. We also calculated sLFO x time in state R-values and quantified the sLFO slope predicting time in state for each scan session and brain state.

### Reproducibility of Mean sLFO x Functional Network Temporal Dynamics

To explore reproducibility of our HCP-YA findings in an independent sample, we followed precisely the same Bayes factor analysis approach as described in the sLFO x DMN-network connectivity analysis.

The first set of eight Bayes factors tested for evidence of reproducibility across sessions, within the HCP-YA sample by comparing strength of the null model (no sLFO x session interaction) to the alternate (sLFO x session interaction). The second set of eight Bayes factors tested for evidence for reproducibility across independent samples, the HCP-YA sample and the healthy adults placebo cohort, by comparing strength of the null model (no sLFO x sample interaction) to the alternate (sLFO x sample interaction). Again, in both cases, Bayes factors which provide evidence for the null model support reproducibility (BF>1), whereas Bayes factors which provide evidence for the alternate provide evidence of a session or sample-based difference in the relationship between sLFO and network temporal dynamics (BF<1).

### Within Subject Changes in Mean sLFO and Functional Brain Network Properties

In addition to testing relationships between the mean sLFO and functional network properties *between* subjects, we also sought to test whether within subject changes in mean sLFO were related to within subject changes in functional brain network properties. We calculated the change in mean sLFO from session 1 to session 2. We also calculated the change in 1) DMN-network connectivity values and 2) percentage of time in brain state from session 1 to session 2. We then ran two linear regression models to test the relationships between 1) ΔsLFO and ΔDMN-network connectivity and 2) ΔsLFO and Δtime in state, which controlled for age, sex, and phase encode direction. Post-hoc analysis included running linear models for each DMN-network / brain state and calculating the R-values for each correlation across each phase encode direction. Post-hoc analysis for DMN-network connectivity also compared the strength of ΔsLFO slopes across networks, mirroring the between subjects approach.

### Replicating Mean sLFO x Brain Relationships with Traditional Arousal Indices

We aimed to confirm that the relationships between mean sLFO and 1) DMN-network connectivity and 2) network temporal dynamics mirrored physiology x brain relationships using more traditional indices. The alternate physiological indices include RVT, HF-HRV, and LF-HRV. In this analysis, 29 subjects were missing physiological data, so our total N=433. The models contained one average measurement indexing physiology for each subject and each scan run, mirroring our sLFO-based models. Where a physiology index x network interaction was significant in predicting connectivity, post-hoc analysis was conducted in the same manner as the sLFO analysis.

### Mean sLFO, Functional Brain Networks, and Catecholaminergic Pharmacology

In our pharmacological manipulation cohort, we tested whether catecholaminergic modulation via pharmacology impacted the relationship between sLFO and 1) DMN-network connectivity and 2) network temporal dynamics through two linear models which tested the effect of sLFO, a sLFO x drug interaction, a sLFO x network interaction, and a three way sLFO x drug x network interaction in relating to connectivity (first linear model) and temporal dynamics (second linear model) while controlling for age, sex, and drug. Following significant sLFO x network interactions and no drug x sLFO x network interactions, we ran linear models for each DMN-network / brain state which controlled for age, sex, drug, and within subject random effects.

Next, we tested whether drug-induced changes in sLFO were related to drug-induced changes in DMN-network connectivity and network temporal dynamics. We tested how change in sLFO (drug – placebo) relate to change in 1) DMN-DAN and DMN-SN connectivity and 2) DMN, DAN, and SN temporal dynamics (drug – placebo) through simple correlation tests. This resulted in ten (5 x 2) correlation tests: sLFO x 5 y-variables (DMN-DAN; DMN-SN; DMN_time_; DAN_time_; SN_time_) x 2 drug conditions (methylphenidate – placebo; haloperidol – placebo).

## Supporting information

Supplemental Results

## Acknowledgments

This research was supported [in part] by the Intramural Research Program of the National Institutes of Health (NIH). The contributions of the NIH author(s) were made as part of their official duties as NIH federal employees, are in compliance with agency policy requirements, and are considered Works of the United States Government. However, the findings and conclusions presented in this paper are those of the author(s) and do not necessarily reflect the views of the NIH or the U.S. Department of Health and Human Services. A portion of this work was presented in a poster in the 2025 Annual Meeting for the College on Problems of Drug Dependence.

## Data Availability

Data collected at the National Institute on Drug Abuse (NIDA) may be made available per request and after obtaining a data transfer agreement. Data from the Human Connectome Project is publicly available on the open access Connectome database (https://db.humanconnectome.org/app/template/Login.vm).

## Code Availability

The code and instructions for performing RIPTiDe denoising using the rapidtide package are publicly available at https://rapidtide.readthedocs.io/en/latest/usage_rapidtide.html#removing-low-frequency-physiological-noise-from-fmri-data.

