## Supplemental Results for "Physiological Arousal as a Predominant Source of Individual Differences in Functional Brain Networks"

### Supplementary Results

Figure S1.

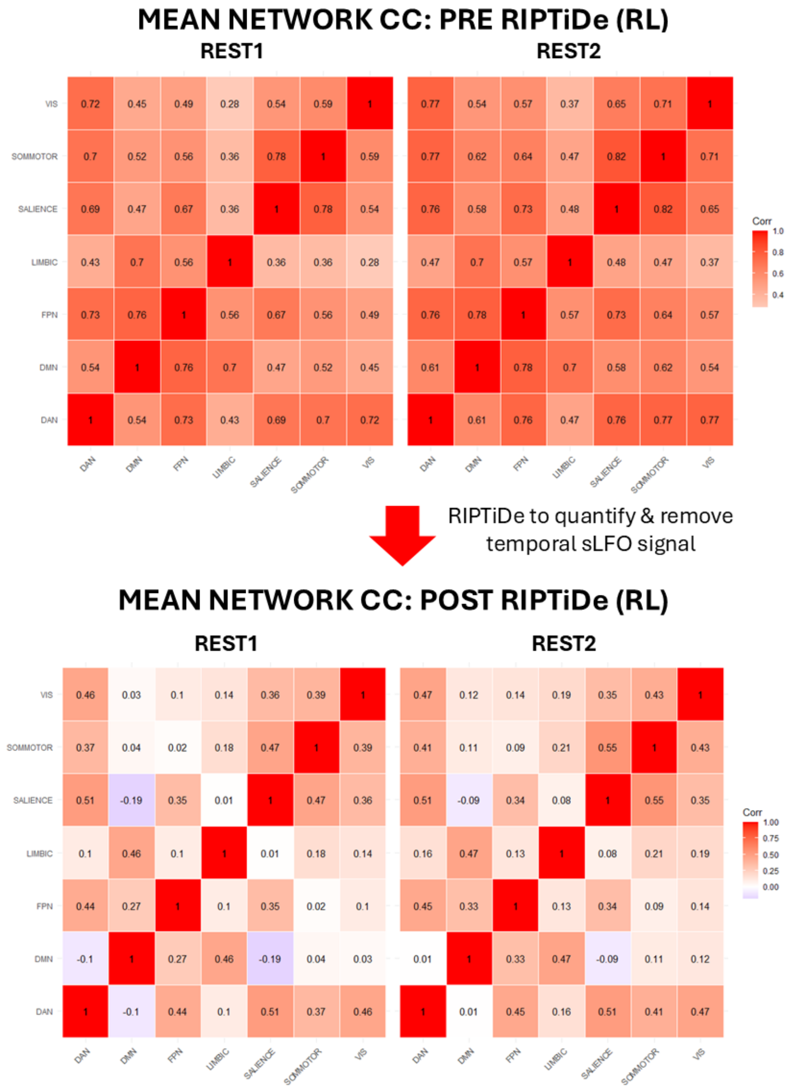

Figure S1. Mean network-network connectivity values across all 7 functional networks before and after RIPTiDe. After RIPTiDe, there is less positive global mean functional connectivity, confirming that regression of sLFO signal using RIPTiDe package reduced global mean functional connectivity.

Table S1.

| Network-network | F | Den df | p (corrected) |
|---|---|---|---|
| **DAN-FPN** | 108.45 | 1830.94 | p<0.001 |
| **DAN-LIM** | 1201.16 | 1805.16 | <0.001 |
| **DAN-SMO** | 1118.79 | 1795.08 | <0.001 |
| **DAN-SN** | 155.82 | 1776.90 | <0.001 |
| **DAN-VIS** | 1131.31 | 1765.98 | <0.001 |
| **FPN-LIM** | 296.34 | 1801.24 | <0.001 |
| **FPN-SMO** | 632.55 | 1779.585 | <0.001 |
| **FPN-SN** | 735.40 | 1832.79 | <0.001 |
| **FPN-VIS** | 551.96 | 1772.865 | <0.001 |
| **LIM-SMO** | 971.065 | 1689.14 | <0.001 |
| **LIM-SN** | 736.015 | 1789.32 | <0.001 |
| **LIM-VIS** | 1043.64 | 1709.30 | <0.001 |
| **SMO-VIS** | 825.12 | 1832.03 | <0.001 |
| **SN-SMO** | 447.035 | 1817.92 | <0.001 |
| **SN-VIS** | 199.98 | 1793.41 | <0.001 |

Table S1. Results for sLFO term in models testing relationship between sLFO and network-network connectivity while controlling for age, sex, session, and phase, for all network-network relationships *except* DMN-network relationships (see Table S2).

Table S2.

| Model Term | num Df | den Df | F | Pr(>F) | NETWORK |
|---|---|---|---|---|---|
| sLFO | 1 | 1817.09 | 1667.98 | <0.001 | DMN-DAN |
| age | 1 | 462.03 | 2.66 | 0.10 | DMN-DAN |
| sex | 1 | 463.17 | 0.30 | 0.58 | DMN-DAN |
| session | 1 | 1383.61 | 2.55 | 0.11 | DMN-DAN |
| phase | 1 | 1383.49 | 1.11 | 0.29 | DMN-DAN |
| sLFO | 1 | 1832.10 | 600.15 | <0.001 | DMN-FPN |
| age | 1 | 460.88 | 1.47 | 0.23 | DMN-FPN |
| sex | 1 | 461.975 | 0.83 | 0.36 | DMN-FPN |
| session | 1 | 1382.54 | 0.35 | 0.55 | DMN-FPN |
| phase | 1 | 1382.43 | 0.46 | 0.50 | DMN-FPN |
| sLFO | 1 | 1727.78 | 206.73 | <0.001 | DMN-LIMBIC |
| age | 1 | 460.03 | 8.63 | 0.0035 | DMN-LIMBIC |
| sex | 1 | 461.28 | 0.92 | 0.34 | DMN-LIMBIC |
| session | 1 | 1381.45 | 0.024 | 0.88 | DMN-LIMBIC |
| phase | 1 | 1381.32 | 1.82 | 0.18 | DMN-LIMBIC |
| sLFO | 1 | 1828.16 | 1360.12 | <0.001 | DMN-SN |
| age | 1 | 461.73 | 0.52 | 0.47 | DMN-SN |
| sex | 1 | 462.84 | 9.72 | 0.0019 | DMN-SN |
| session | 1 | 1383.37 | 1.09 | 0.30 | DMN-SN |
| phase | 1 | 1383.25 | 1.765 | 0.18 | DMN-SN |
| sLFO | 1 | 1816.14 | 1286.27 | <0.001 | DMN-SMN |
| age | 1 | 462.07 | 0.62 | 0.43 | DMN-SMN |
| sex | 1 | 463.22 | 3.79 | 0.052 | DMN-SMN |
| session | 1 | 1383.655 | 2.09 | 0.15 | DMN-SMN |
| phase | 1 | 1383.53 | 0.31 | 0.58 | DMN-SMN |
| sLFO | 1 | 1793.46 | 1351.82 | <0.001 | DMN-VIS |
| age | 1 | 461.85 | 2.65 | 0.10 | DMN-VIS |
| sex | 1 | 463.035 | 7.06 | 0.0082 | DMN-VIS |
| session | 1 | 1383.37 | 6.53 | 0.011 | DMN-VIS |
| phase | 1 | 1383.243 | 1.49 | 0.22 | DMN-VIS |

Table S2. Each DMN-Network model testing relationship between sLFO and network-network connectivity while controlling for age, sex, session, and phase.

Table S3.

| DMN-Network Contrast | estimate | z.ratio | p |
|---|---|---|---|
| DMN-DAN — DMN-FPN | 0.71 | 14.39 | <0.001 |
| DMN-DAN — DMN-LIMBIC | 1.14 | 23.16 | <0.001 |
| DMN-DAN — DMN-SALIENCE | -0.02 | -0.41 | 0.998452 |
| DMN-DAN — DMN-SOMMOTOR | 0.28 | 5.67 | <0.001 |
| DMN-DAN — DMN-VIS | 0.34 | 6.82 | <0.001 |
| DMN-FPN — DMN-LIMBIC | 0.43 | 8.77 | <0.001 |
| DMN-FPN — DMN-SALIENCE | -0.73 | -14.80 | <0.001 |
| DMN-FPN — DMN-SOMMOTOR | -0.43 | -8.72 | <0.001 |
| DMN-FPN — VIS | -0.37 | -7.57 | <0.001 |
| DMN-LIMBIC — DMN-SALIENCE | -1.16 | -23.57 | <0.001 |
| DMN-LIMBIC — DMN-SOMMOTOR | -0.86 | -17.49 | <0.001 |
| DMN-LIMBIC — DMN-VIS | -0.805 | -16.34 | <0.001 |
| DMN-SALIENCE — DMN-SOMMOTOR | 0.300 | 6.09 | <0.001 |
| DMN-SALIENCE — DMN-VIS | 0.36 | 7.24 | <0.001 |
| DMN-SOMMOTOR — DMN-VIS | 0.057 | 1.15 | 0.860397 |

Table S3. Pairwise comparisons between the sLFO slopes for each DMN-network relationship collapsed across session 1 and session 2. Estimate reflects the difference in sLFO slope between the two DMN-network connectivity models. Models controlled for age, sex, phase-encoding direction and session. SE=0.049 and is consistent for each comparison because it is calculated as a result of the whole model which includes each network-network value. Degrees of freedom are asymptotic (approach infinity).

Table S4.

| Model Term | num Df | den Df | F | p | state |
|---|---|---|---|---|---|
| sLFO | 1 | 1769.93 | 697.40 | <0.001 | FI-DMN |
| age | 1 | 461.63 | 27.10 | <0.001 | FI-DMN |
| sex | 1 | 462.84 | 25.12 | <0.001 | FI-DMN |
| phase | 1 | 922.80 | 0.63 | 0.43 | FI-DMN |
| session | 1 | 460.67 | 0.10 | 0.755 | FI-DMN |
| sLFO | 1 | 1712.70 | 7.28 | 0.007 | FPN |
| age | 1 | 462.04 | 2.52 | 0.11 | FPN |
| sex | 1 | 463.30 | 2.49 | 0.115 | FPN |
| phase | 1 | 921.90 | 0.06 | 0.81 | FPN |
| session | 1 | 460.87 | 38.35 | <0.001 | FPN |
| sLFO | 1 | 1786.16 | 1016.13 | <0.001 | DMN |
| age | 1 | 461.70 | 0.63 | 0.43 | DMN |
| sex | 1 | 462.88 | 4.01 | 0.046 | DMN |
| phase | 1 | 922.83 | 0.56 | 0.45 | DMN |
| session | 1 | 460.75 | 2.22 | 0.14 | DMN |
| sLFO | 1 | 1829.68 | 546.99 | <0.001 | DAN |
| age | 1 | 461.08 | 7.75 | 0.0056 | DAN |
| sex | 1 | 462.18 | 5.04 | 0.025 | DAN |
| phase | 1 | 920.92 | 1.78 | 0.18 | DAN |
| session | 1 | 460.86 | 16.10 | <0.001 | DAN |
| sLFO | 1 | 1617.47 | 626.55 | <0.001 | SN-1 |
| age | 1 | 462.01 | 4.66 | 0.031 | SN-1 |
| sex | 1 | 463.31 | 0.14 | 0.71 | SN-1 |
| phase | 1 | 923.06 | 0.00 | 0.98 | SN-1 |
| session | 1 | 462.02 | 11.27 | 0.00085 | SN-1 |
| sLFO | 1 | 1698.49 | 824.27 | <0.001 | SMO |
| age | 1 | 462.38 | 34.79 | <0.001 | SMO |
| sex | 1 | 463.65 | 20.85 | <0.001 | SMO |
| phase | 1 | 922.57 | 1.59 | 0.21 | SMO |
| session | 1 | 460.64 | 0.06 | 0.81 | SMO |
| sLFO | 1 | 1752.31 | 7.61 | 0.00585 | SM-DMN |
| age | 1 | 461.27 | 10.07 | 0.0016 | SM-DMN |
| sex | 1 | 462.49 | 11.01 | <0.001 | SM-DMN |
| phase | 1 | 922.49 | 0.73 | 0.39 | SM-DMN |
| session | 1 | 460.90 | 2.48 | 0.12 | SM-DMN |
| sLFO | 1 | 1786.22 | 0.11 | 0.74 | SN-2 |
| age | 1 | 460.40 | 47.49 | <0.001 | SN-2 |
| sex | 1 | 461.58 | 22.82 | <0.001 | SN-2 |

| | | | | | |
|---|---|---|---|---|---|
| phase | 1 | 922.53 | 0.00 | 0.97 | SN-2 |
| session | 1 | 460.44 | 25.39 | <0.001 | SN-2 |

Table S4. Each brain state model testing relationship between sLFO and network temporal dynamics while controlling for age, sex, session, and phase.

#### Investigating age-related effects on the relationship between sLFO and brain state temporal dynamics

There was a significant age x sLFO x brain state interaction in the HCP-YA sample, in a model predicting percentage of time in state ($F=4.06$, $DOF=7/14121.23$, $p<0.001$). After correcting for eight multiple comparisons, only interactions with SN-1 ($F=8.47$, $DOF=1/1617.69$, $p_{corr}=0.02$) and DAN ($F=7.335$, $DOF=1/1829.47$, $p_{corr}=0.054$) survived, with DAN at trend-level. For both DAN and SN-1, there was a stronger relationship between sLFO and percent of time in brain network state in younger individuals compared to older. Before multiple comparisons corrections, there were also trend-level sLFO x age interactions for the FI-DMN ($F=3.2$, $DOF=1/1766.95$, $p_{raw}=0.07$) and FPN ($F=3.2$, $DOF=1/1718.37$, $p_{raw}=0.07$) states.

Figure S2.

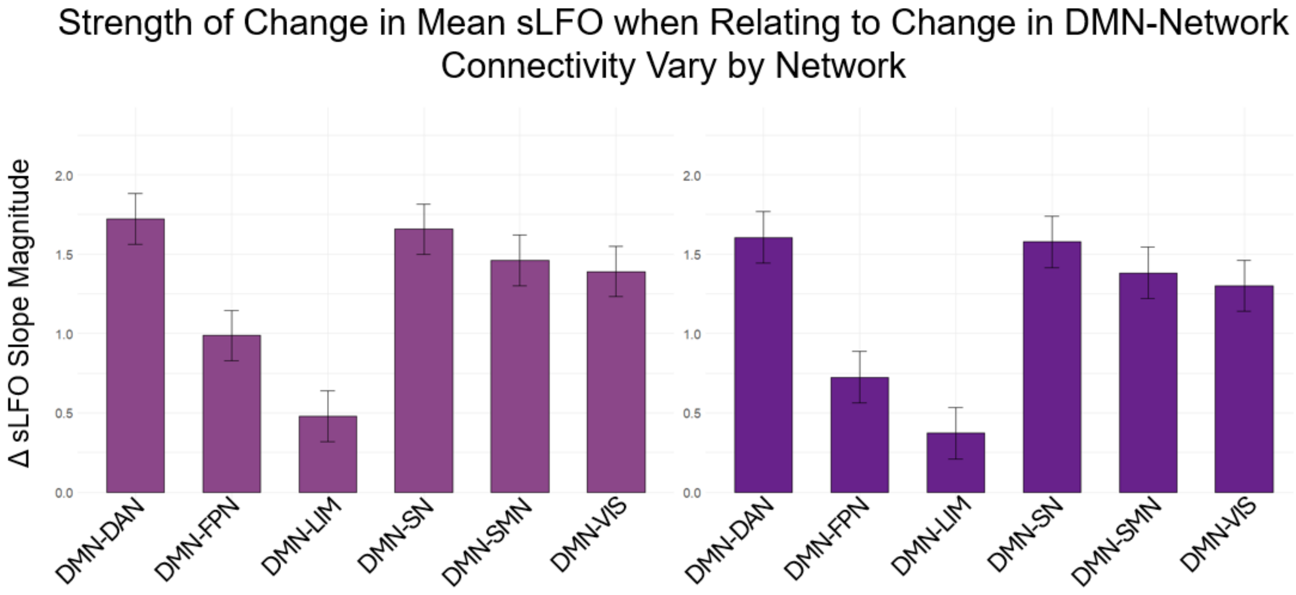

Figure S2. Magnitude of change in sLFO slopes (REST1 – REST2) predicting change in network-network connectivity (REST1-REST2). First 6 network-network values are data from LR phase encoding. Second 6 network-network values are data from RL phase encoding. Error bars represent upper/lower limits of trend prediction.

Table S5.

| **DMN-Network Contrast** | **estimate** | **z.ratio** | **p.value** |
|---|---|---|---|
| DMN-DAN – DMN-FPN | 0.83 | 12.175 | <0.001 |
| DMN-DAN – LIMBIC | 1.33 | 19.52 | <0.001 |
| DMN-DAN – DMN-SALIENCE | 0.05 | 0.75 | 0.98 |
| DMN-DAN – DMN-SOMMOTOR | 0.31 | 4.60 | <0.001 |
| DMN-DAN – DMN-VIS | 0.34 | 4.95 | <0.001 |
| DMN-FPN – DMN-LIMBIC | 0.50 | 7.34 | <0.001 |
| DMN-FPN – DMN-SALIENCE | -0.78 | -11.43 | <0.001 |
| DMN-FPN – DMN-SOMMOTOR | -0.52 | -7.57 | <0.001 |
| DMN-FPN – DMN-VIS | -0.49 | -7.23 | <0.001 |
| DMN-LIMBIC – DMN-SALIENCE | -1.28 | -18.77 | <0.001 |
| DMN-LIMBIC – DMN-SOMMOTOR | -1.02 | -14.92 | <0.001 |
| DMN-LIMBIC – DMN-VIS | -0.99 | -14.57 | <0.001 |
| DMN-SALIENCE – DMN-SOMMOTOR | 0.26 | 3.85 | 0.0016 |
| DMN-SALIENCE – DMN-VIS | 0.29 | 4.20 | <0.001 |
| DMN-SOMMOTOR – DMN-VIS | 0.0235 | 0.345 | 0.999 |

Table S5. Pairwise comparisons between the change in sLFO slope (REST1-REST2) and change in each DMN-network relationship (REST1-REST2). Models controlled for age, sex, and phase encoding. SE=0.07 and is consistent for each comparison because it is calculated as a result of the whole model which includes each network-network value. Degrees of freedom are asymptotic (approach infinity).

Table S6.

| Model Term | den Df | F | p | state |
|---|---|---|---|---|
| sLFO (REST2 - REST1) | 1843.00 | 535.73 | <0.001 | FI-DMN |
| age | 459.00 | 0.16 | 0.69 | FI-DMN |
| sex | 458.92 | 0.80 | 0.37 | FI-DMN |
| phase | 1470.29 | 44.33 | <0.001 | FI-DMN |
| sLFO (REST2 - REST1) | 1832.56 | 1.04 | 0.31 | FPN |
| age | 457.96 | 0.01 | 0.93 | FPN |
| sex | 457.87 | 2.16 | 0.14 | FPN |
| phase | 1477.08 | 0.00 | 0.99 | FPN |
| sLFO (REST2 - REST1) | 1833.34 | 880.56 | <0.001 | DMN |
| age | 458.52 | 0.03 | 0.87 | DMN |
| sex | 458.43 | 0.14 | 0.71 | DMN |
| phase | 1477.29 | 63.40 | <0.001 | DMN |
| sLFO (REST2 - REST1) | 1842.06 | 560.60 | <0.001 | DAN |
| age | 458.52 | 2.18 | 0.14 | DAN |
| sex | 458.44 | 0.03 | 0.86 | DAN |
| phase | 1472.37 | 50.59 | <0.001 | DAN |
| sLFO (REST2 - REST1) | 1804.79 | 547.83 | <0.001 | SN-1 |
| age | 458.02 | 1.45 | 0.23 | SN-1 |
| sex | 457.93 | 2.01 | 0.16 | SN-1 |
| phase | 1483.31 | 24.08 | <0.001 | SN-1 |
| sLFO (REST2 - REST1) | 1841.81 | 733.04 | <0.001 | SMN |
| age | 458.29 | 0.36 | 0.55 | SMN |
| sex | 458.21 | 3.56 | 0.06 | SMN |
| phase | 1467.09 | 72.34 | <0.001 | SMN |
| sLFO (REST2 - REST1) | 1824.77 | 9.14 | 0.0025 | SM-DMN |
| age | 458.64 | 0.01 | 0.92 | SM-DMN |
| sex | 458.55 | 0.13 | 0.72 | SM-DMN |
| phase | 1479.92 | 27.49 | <0.001 | SM-DMN |
| sLFO (REST2 - REST1) | 1842.97 | 1.87 | 0.17 | SN-2 |
| age | 459.01 | 0.04 | 0.84 | SN-2 |
| sex | 458.93 | 3.97 | 0.047 | SN-2 |
| phase | 1470.03 | 40.43 | <0.001 | SN-2 |

Table S6. Within subject F, DOF and p-value for models predicting change in percent time in state from change in sLFO from REST1 to REST2. Num df = 1 for all models/terms. Models controlled for sex, age, and phase.

Figure S3.

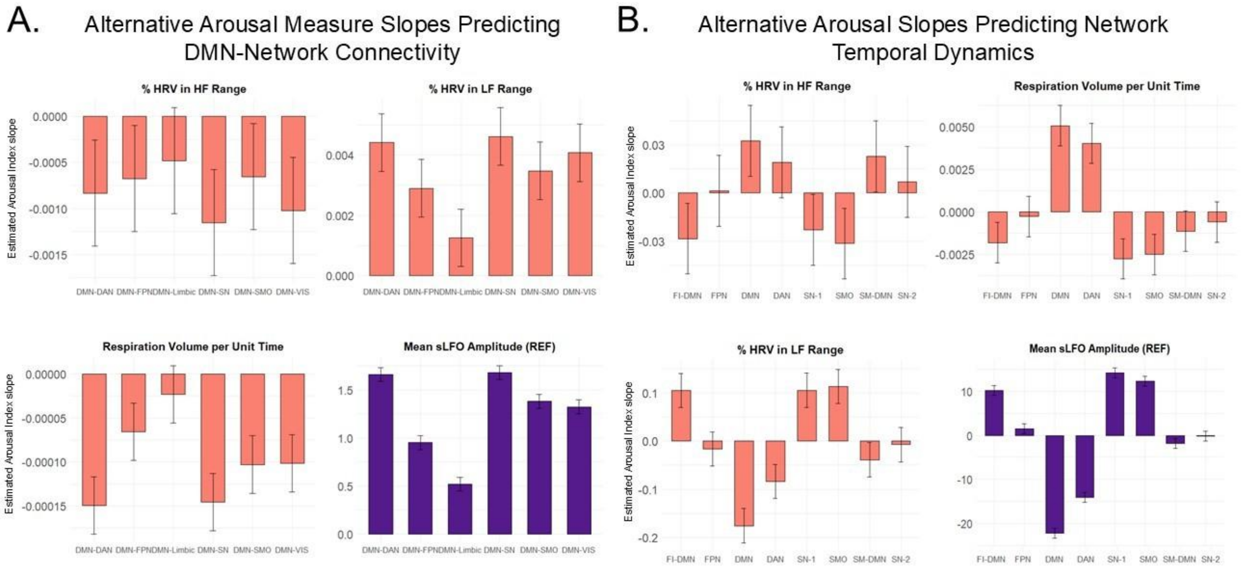

Figure S3. A. Alternative arousal measure slopes predicting DMN-network connectivity by network. B. Alternative arousal slopes predicting network temporal dynamics, by brain state. For both plots, orange indicates alternate arousal measure; purple indicates sLFO slope for reference. Alternative arousal measures include HF-HRV, LF-HRV, and RVT. RVT: Respiration Volume per Unit Time; HF-HRV: Percent of heart rate variability in high frequency range; LF-HRV: Percent of heart rate variability in low frequency range.

Table S7

| Model term | num Df | den Df | F | p |
|---|---|---|---|---|
| RVT | 1 | 7915.26 | 136.24 | <0.001 |
| network | 5 | 2145.11 | 2134.62 | <0.001 |
| session | 1 | 7561.98 | 2.31 | 0.128598 |
| phase | 1 | 7534.77 | 0.38 | 0.536845 |
| sex | 1 | 426.70 | 10.58 | 0.0012 |
| age | 1 | 427.30 | 3.67 | 0.056 |
| network x RVT | 5 | 7360.72 | 9.40 | <0.001 |
| HF-HRV | 1 | 7513.89 | 23.31 | <0.001 |
| network | 5 | 2144.49 | 2109.73 | <0.001 |
| session | 1 | 7569.11 | 1.16 | 0.28 |
| phase | 1 | 7534.47 | 0.06 | 0.80 |
| sex | 1 | 438.65 | 6.84 | 0.0092 |
| age | 1 | 426.32 | 1.77 | 0.185 |
| network x HF-HRV | 5 | 4990.86 | 0.90 | 0.48 |
| LF-HRV | 1 | 7840.69 | 182.56 | <0.001 |
| network | 5 | 2144.53 | 2117.68 | <0.001 |
| session | 1 | 7560.41 | 1.32 | 0.25 |
| phase | 1 | 7533.74 | 0.00 | 0.97 |
| sex | 1 | 430.91 | 5.02 | 0.0255 |
| age | 1 | 426.35 | 1.54 | 0.215 |
| network x LF-HRV | 5 | 6352.69 | 7.60 | <0.001 |

Table S7. Three model results for each alternate indices of arousal relating to DMN-network connectivity. All physiology measures (RVT, HF-HRV, LF-HRV) were demeaned in each model. RVT: respiration volume per unit time; HF-HRV: Percent of heart rate variability in high frequency range; LF-HRV: Percent of heart rate variability in low frequency range.

Table S8.

| INDEX | contrast | estimate | z.ratio | p.value | pcor |
|---|---|---|---|---|---|
| | DAN - FPN | -8.39E-05 | -3.77 | 0.0022 | 0.033462 |
| | DAN - LIMBIC | -0.00013 | -5.68 | <0.001 | 2.95E-06 |
| | DAN - SALIENCE | -3.88E-06 | -0.17 | 1 | 1 |
| | DAN - SOMMOTOR | -4.64E-05 | -2.09 | 0.29 | 1 |
| | DAN - VIS | -4.76E-05 | -2.14 | 0.27 | 1 |
| | FPN - LIMBIC | -4.25E-05 | -1.91 | 0.395 | 1 |
| | FPN - SALIENCE | 8.00E-05 | 3.60 | 0.004 | 0.065 |
| | FPN - SOMMOTOR | 3.75E-05 | 1.69 | 0.54 | 1 |
| | FPN - VIS | 3.63E-05 | 1.63 | 0.58 | 1 |
| | LIMBIC - SALIENCE | 0.00012 | 5.51 | <0.001 | <0.001 |
| | LIMBIC - SOMMOTOR | 8.00E-05 | 3.60 | 0.0044 | 0.0655 |
| | LIMBIC - VIS | 7.88E-05 | 3.54 | 0.0053 | 0.079 |
| | SALIENCE - SOMMOTOR | -4.26E-05 | -1.91 | 0.39 | 1 |
| | SALIENCE - VIS | -4.37E-05 | -1.97 | 0.36 | 1 |
| **RVT** | SOMMOTOR - VIS | -1.16E-06 | -0.05 | 1 | 1 |
| | DAN - FPN | 0.0015 | 2.36 | 0.17 | 1 |
| | DAN - LIMBIC | 0.0032 | 4.93 | <0.001 | <0.001 |
| | DAN - SALIENCE | -0.0002 | -0.31 | 1 | 1 |
| | DAN - SOMMOTOR | 0.00094 | 1.47 | 0.68 | 1 |
| | DAN - VIS | 0.00034 | 0.53 | 0.995 | 1 |
| | FPN - LIMBIC | 0.0016 | 2.57 | 0.11 | 1 |
| | FPN - SALIENCE | -0.0017 | -2.68 | 0.08 | 1 |
| | FPN - SOMMOTOR | -0.00057 | -0.89 | 0.95 | 1 |
| | FPN - VIS | -0.0012 | -1.83 | 0.445 | 1 |
| | LIMBIC - SALIENCE | -0.0034 | -5.24 | <0.001 | <0.001 |
| | LIMBIC - SOMMOTOR | -0.0022 | -3.46 | 0.00715 | 0.11 |
| | LIMBIC - VIS | -0.0028 | -4.40 | <0.001 | 0.0024 |
| | SALIENCE - SOMMOTOR | 0.0011 | 1.78 | 0.48 | 1 |
| | SALIENCE - VIS | 0.00054 | 0.84 | 0.96 | 1 |
| **LF-HRV** | SOMMOTOR - VIS | -0.0006 | -0.94 | 0.94 | 1 |

Table S8. Pairwise comparisons of strength of the slope of arousal indices predicting network-network connectivity. RVT: respiration volume per unit time; LF-HRV: Percent of heart rate variability in low frequency range.

Table S9

| Model term | num Df | den Df | F | p |
|---|---|---|---|---|
| RVT | 1 | 13453.62 | 3.48E-05 | 1 |
| brain state | 7 | 3417.26 | 3184.787 | 0 |
| session | 1 | 10217.15 | 5.94E-06 | 1 |
| phase | 1 | 10091.25 | 2.84E-06 | 1 |
| sex | 1 | 3420.02 | 3.10E-06 | 1 |
| age | 1 | 3438.07 | 1.72E-05 | 1 |
| brain state x RVT | 7 | 13456.6 | 23.76 | <0.001 |
| brain state x sex | 7 | 3420.39 | 14.45 | <0.001 |
| brain state x age | 7 | 3438.48 | 12.18 | <0.001 |
| HF-HRV | 1 | 12222.63 | 2.64E-06 | 1 |
| brain state | 7 | 3420.29 | 3110.18 | <0.001 |
| session | 1 | 10233.52 | 4.45E-06 | 1 |
| phase | 1 | 10078.81 | 3.80E-06 | 1 |
| sex | 1 | 3670.76 | 2.61E-06 | 1 |
| age | 1 | 3415.26 | 9.08E-06 | 1 |
| brain state x HF-HRV | 7 | 12246.98 | 4.88 | <0.001 |
| brain state x sex | 7 | 3670.54 | 10.37 | <0.001 |
| brain state x age | 7 | 3415.63 | 11.18 | <0.001 |
| LF-HRV | 1 | 13192.01 | 7.11E-06 | 1 |
| brain state | 7 | 3419.75 | 3158.89 | <0.001 |
| session | 1 | 10212.32 | 5.17E-06 | 1 |
| phase | 1 | 10086.39 | 4.01E-06 | 1 |
| sex | 1 | 3531.55 | 7.05E-07 | 1 |
| age | 1 | 3416.20 | 1.03E-05 | 1 |
| brain state x LF-HRV | 7 | 13196.18 | 32.64 | <0.001 |
| brain state x sex | 7 | 3531.93 | 9.79 | <0.001 |
| brain state x age | 7 | 3416.66 | 11.22 | <0.001 |

Table S9. Three model results for each alternate indices of arousal in predicting network temporal dynamics. All physiology measures (RVT, HF-HRV, LF-HRV) and age were demeaned. RVT: respiration volume per unit time; HF-HRV: Percent of heart rate variability in high frequency range; LF-HRV: Percent of heart rate variability in low frequency range.

Table S10.

| Model Term | num Df | den Df | F | p |
|---|---|---|---|---|
| Drug | 2 | 695.48 | 4.17 | 0.016 |
| Network | 5 | 321.59 | 173.07 | p<0.001 |
| sLFO | 1 | 668.94 | 119.45 | p<0.001 |
| age | 1 | 56.17 | 5.65 | 0.021 |
| Sex | 1 | 56.35 | 0.67 | 0.415 |
| Drug:Network | 10 | 726.46 | 1.25 | 0.255 |
| Drug:sLFO | 2 | 715.64 | 2.93 | 0.054 |
| Network:sLFO | 5 | 790.39 | 5.865 | p<0.001 |
| Drug:Network:sLFO | 10 | 899.99 | 0.88 | 0.55 |

Table S10. Pharmacological manipulation sample DMN-network main model results for DMN x 6 functional networks. Den df approximated via S-method. Network term includes: DMN-FPN, DMN-Limbic, DMN-DAN, DMN-SN, DMN-SMO, and DMN-VIS. sLFO was demeaned in the model.

Table S11.

| Model Term | num Df | den Df | F | Pr(>F) | NETWORK |
|---|---|---|---|---|---|
| sLFO | 1 | 168.98 | 39.36 | <0.001 | DMN-DAN |
| drug | 2 | 121.72 | 0.91 | 0.41 | DMN-DAN |
| age | 1 | 55.90 | 4.28 | 0.04 | DMN-DAN |
| sex | 1 | 56.23 | 0.64 | 0.43 | DMN-DAN |
| sLFO | 1 | 170.22 | 51.70 | <0.001 | DMN-FPN |
| drug | 2 | 121.22 | 1.64 | 0.20 | DMN-FPN |
| age | 1 | 55.57 | 3.35 | 0.07 | DMN-FPN |
| sex | 1 | 55.90 | 0.12 | 0.73 | DMN-FPN |
| sLFO | 1 | 170.98 | 4.52 | 0.03 | DMN-LIMBIC |
| drug | 2 | 121.18 | 0.39 | 0.68 | DMN-LIMBIC |
| age | 1 | 55.98 | 2.18 | 0.15 | DMN-LIMBIC |
| sex | 1 | 56.29 | 0.39 | 0.54 | DMN-LIMBIC |
| sLFO | 1 | 169.98 | 49.15 | <0.001 | DMN-SN |
| drug | 2 | 121.13 | 1.34 | 0.27 | DMN-SN |
| age | 1 | 55.43 | 6.30 | 0.02 | DMN-SN |
| sex | 1 | 55.76 | 0.70 | 0.41 | DMN-SN |
| sLFO | 1 | 171.00 | 25.58 | <0.001 | DMN-SMN |
| drug | 2 | 121.19 | 2.05 | 0.13 | DMN-SMN |
| age | 1 | 55.91 | 3.46 | 0.07 | DMN-SMN |
| sex | 1 | 56.23 | 1.40 | 0.24 | DMN-SMN |
| sLFO | 1 | 145.33 | 13.06 | <0.001 | DMN-VIS |
| drug | 2 | 119.56 | 5.38 | 0.01 | DMN-VIS |
| age | 1 | 52.74 | 8.82 | 0.004 | DMN-VIS |
| sex | 1 | 53.08 | 2.81 | 0.10 | DMN-VIS |

Table S11. Each network-network connectivity by sLFO model controlling for drug, age, and sex with raw uncorrected p values.

Table S12

| Model Term | num Df | den Df | F | p |
|---|---|---|---|---|
| sLFO | 1 | 1235.96 | 2.71E-12 | 1 |
| brain state | 7 | 479.76 | 296.7 | p<0.001 |
| Drug | 2 | 970.60 | 3.10E-29 | 1 |
| sLFO:brain state | 7 | 1235.96 | 28.875 | p<0.001 |
| sLFO:Drug | 2 | 1174.73 | 1.21E-30 | 1 |
| brain state:Drug | 14 | 970.60 | 3.74 | p<0.001 |
| brain state:age | 8 | 443.90 | 1.97 | 0.049 |
| brain state:Sex | 8 | 446.9 | 0.44 | 0.90 |
| sLFO:brain state:Drug | 14 | 1174.73 | 1.48 | 0.11 |

Table S12. Pharmacological manipulation sample main model results for percentage time in eight brain states. Den df approximated via S-method. Brain state term includes: FI-DMN, FPN, DMN, DAN, SN-1, SMO, SM-DMN, SN-2. Both sLFO and age were demeaned.

Table S13.

| Brain state | variable | num Df | den Df | F | p | p (corrected) |
|---|---|---|---|---|---|---|
| **FI-DMN** | sLFO | 1 | 158.77 | 1.81 | 0.18 | 1 |
| | Drug | 2 | 122.53 | 0.52 | 0.59 | |
| | Age | 1 | 55.88 | 3.17 | 0.08 | |
| | Sex | 1 | 56.23 | 0.67 | 0.42 | |
| **FPN** | sLFO | 1 | 150.35 | 33.35 | <0.001 | <0.001 |
| | Drug | 2 | 121.74 | 3.58 | 0.03 | |
| | Age | 1 | 54.82 | 0.03 | 0.88 | |
| | Sex | 1 | 55.17 | 0.00 | 0.98 | |
| **DMN** | sLFO | 1 | 156.87 | 52.61 | <0.001 | <0.001 |
| | Drug | 2 | 121.62 | 4.30 | 0.02 | |
| | Age | 1 | 54.85 | 0.07 | 0.79 | |
| | Sex | 1 | 55.21 | 0.19 | 0.67 | |
| **DAN** | sLFO | 1 | 162.61 | 42.42 | <0.001 | <0.001 |
| | Drug | 2 | 121.08 | 4.47 | 0.01 | |
| | Age | 1 | 54.52 | 0.90 | 0.35 | |
| | Sex | 1 | 54.86 | 0.04 | 0.85 | |
| **SN-1** | sLFO | 1 | 163.71 | 20.50 | <0.001 | <0.001 |
| | Drug | 2 | 122.29 | 1.35 | 0.26 | |
| | Age | 1 | 55.86 | 0.42 | 0.52 | |
| | Sex | 1 | 56.20 | 0.62 | 0.43 | |
| **SMO** | sLFO | 1 | 164.70 | 0.73 | 0.40 | 1 |
| | Drug | 2 | 122.11 | 1.73 | 0.18 | |
| | Age | 1 | 55.72 | 2.78 | 0.10 | |
| | Sex | 1 | 56.07 | 0.06 | 0.81 | |
| **SM-DMN** | sLFO | 1 | 134.79 | 2.58 | 0.11 | 0.884 |
| | Drug | 2 | 121.98 | 5.55 | 0.00 | |
| | Age | 1 | 54.93 | 3.89 | 0.05 | |
| | Sex | 1 | 55.27 | 1.76 | 0.19 | |
| **SN-2** | sLFO | 1 | 162.62 | 16.58 | <0.001 | <0.001 |
| | Drug | 2 | 122.23 | 2.00 | 0.14 | |
| | Age | 1 | 55.73 | 7.23 | 0.01 | |
| | Sex | 1 | 56.08 | 0.34 | 0.56 | |

Table S13. Each brain state model testing relationship between sLFO and percentage of time in state while controlling for drug, age, and sex. Raw uncorrected p values provided for all terms and Bonferroni corrected p values provided for sLFO term (N comparisons = 8).

Figure S4.

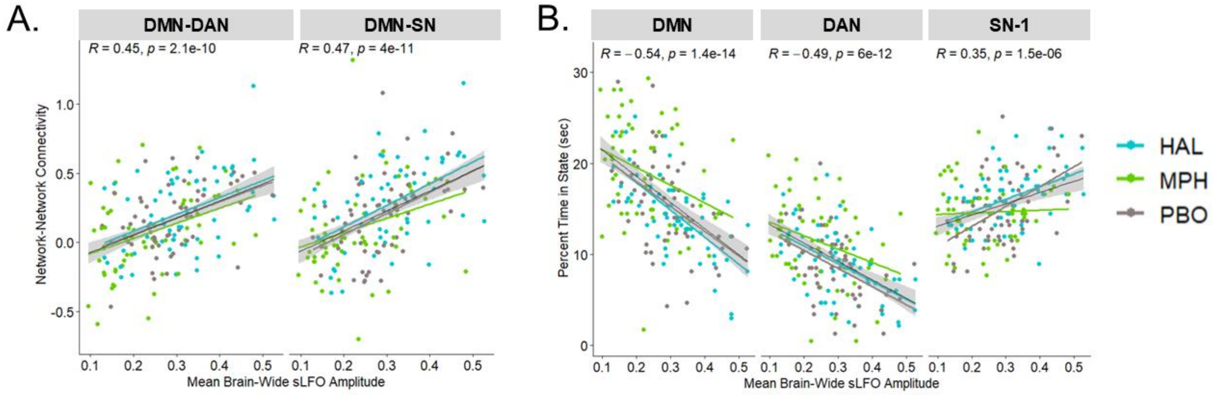

Figure S4. A. Plots of the relationship between mean sLFO amplitude and DMN-DAN and DMN-SN connectivity, under each pharmacological condition. B. Plots of the relationship between mean sLFO amplitude and time spent in the DMN, DAN, and SN, under each pharmacological condition. DMN: Default Mode Network; DAN: Dorsal Attention Network; SN: Salience Network; HAL: haloperidol; MPH: methylphenidate; PBO: placebo.
